# Population genomics reveals strong impacts of genetic drift without purging and guides conservation of bull and giant kelp

**DOI:** 10.1101/2024.10.10.617648

**Authors:** Jordan B. Bemmels, Samuel Starko, Brooke L. Weigel, Kaede Hirabayashi, Alex Pinch, Cassandra Elphinstone, Megan N. Dethier, Loren H. Rieseberg, Jonathan E. Page, Christopher J. Neufeld, Gregory L. Owens

## Abstract

Kelp forests are declining in many parts of the northeast Pacific^1–4^. In small populations, genetic drift can reduce adaptive variation and increase fixation of recessive deleterious alleles^5–7^, but natural selection may purge harmful variants^8–10^. To understand evolutionary dynamics and inform restoration strategies, we investigated genetic structure and the outcomes of genetic drift and purging by sequencing the genomes of 429 bull kelp (*Nereocystis luetkeana*) and 211 giant kelp (*Macrocystis pyrifera* sensu lato^11^; cf. ^12^) from the coastlines of British Columbia and Washington. We identified 6-7 geographically and genetically distinct clusters in each species. Low effective population size was associated with low genetic diversity and high inbreeding coefficients (including increased selfing rates), with extreme variation in these genetic health indices among bull kelp populations but more moderate variation in giant kelp. We found no evidence that natural selection is purging putative recessive deleterious alleles in either species. Instead, genetic drift has fixed many such alleles in small populations of bull kelp, leading us to predict (1) reduced within-population inbreeding depression in small populations, which may be associated with an observed shift toward increased selfing rate, and (2) hybrid vigour in crosses between small populations. Our genomic findings imply several strategies for optimal sourcing and crossing of populations for restoration and aquaculture, but which require experimental validation. Overall, our work reveals strong genetic structure and suggests that conservation strategies should consider the multiple health risks faced by small populations whose evolutionary dynamics are dominated by genetic drift.

## Results and discussion

Bull kelp and giant kelp are the principal canopy-forming species in kelp forests of the northeast Pacific, supporting highly productive and biodiverse ecosystems^13^ that generate billions of dollars annually in ecosystem services^14^. Despite their broad geographic distributions from Alaska to California (with giant kelp additionally found in northern Mexico and the Southern Hemisphere^13^) both species have experienced strong local and regional declines^1–4^ due to factors such as marine heatwaves and urchin overgrazing^2,3^. These declines have spurred a profusion of interest in kelp forest restoration^15,16^. However, restoration and conservation strategies for bull and giant kelp are being hampered in part by lack of information about genetic structure (cf. ^17,18^), patterns of local adaptation, and the genetic risks faced by small populations subject to the balance between genetic drift and natural selection. To address this knowledge gap, we sequenced the genomes of 429 bull kelp and 211 giant kelp (Table S1) from British Columbia (BC), Canada and Washington (WA), USA (hereafter, ‘BCWA’; Figure S1) to a mean depth of 21.0X (range: 11.5-43.0X). We identified 3,274,934 autosomal single nucleotide polymorphisms (SNPs) with a minimum minor allele frequency of 0.01 in bull kelp and 2,341,413 such SNPs in giant kelp, or 4,327,335 SNPs when including published data^19–21^ from an additional 70 individuals from California, Chile, and Australia. Southern and Northern Hemisphere giant kelp formed distinct genetic clusters^22^ (Figure S2A,C) and were distinguished along the first principal component (PC) axis of genetic variation (Figure S2E). The hemispheres were highly genetically differentiated (*F*_ST_ = 0.71) and moderately genetically diverged (*d*_XY_ = 0.0077). Within North America there are two giant kelp ecomorphs (*integrifolia* and *pyrifera*) whose geographic distributions partly overlap in California^11,19^. After excluding close relatives (Table S1) only one giant kelp sample from California was of the *integrifolia* ecomorph, preventing calculation of population statistics. The remaining 10 Californian *pyrifera* individuals were moderately genetically distinct from BCWA (all *integrifolia*^11,19^) (*F*_ST_ = 0.34; *d*_XY_ = 0.0040; Figure S2). Due to geographic, genetic and ecomorph differences, the remainder of our analyses will focus on BCWA only.

Within BCWA, both species exhibit strong genetic structure (Table S2). Six genetic clusters were identified in bull kelp (Figure 1A,C) and seven clusters in giant kelp (Figure 1B,D) (see also Figure S3). These clusters occupy distinct geographic regions and are largely non-overlapping along the first two PC axes of genetic variation (Figure 1E-F). A strong isolation-by-distance pattern of increasing genetic distance (*d*_XY_) with geographic distance (Figure S4) and the presence of populations admixed between clusters (Figure 1A-D) suggest that adjacent clusters are connected by gene flow. Given the lack of formal kelp management zones in BCWA, guidelines are needed to inform movement of genetic material for restoration and aquaculture. We suggest that genetic clusters could be used to help define Management Units (MUs)^23,24^ or – in combination with environmental data^25^ – seed-transfer zones^26^ that would delineate regions within which transfer of genetic material would pose minimal risk to the genetic integrity of local populations. Moreover, genetic clusters can be used to guide biobanking efforts and prioritize conservation investments, given that small populations in some clusters may be at risk for extirpation.

**Figure 1.**
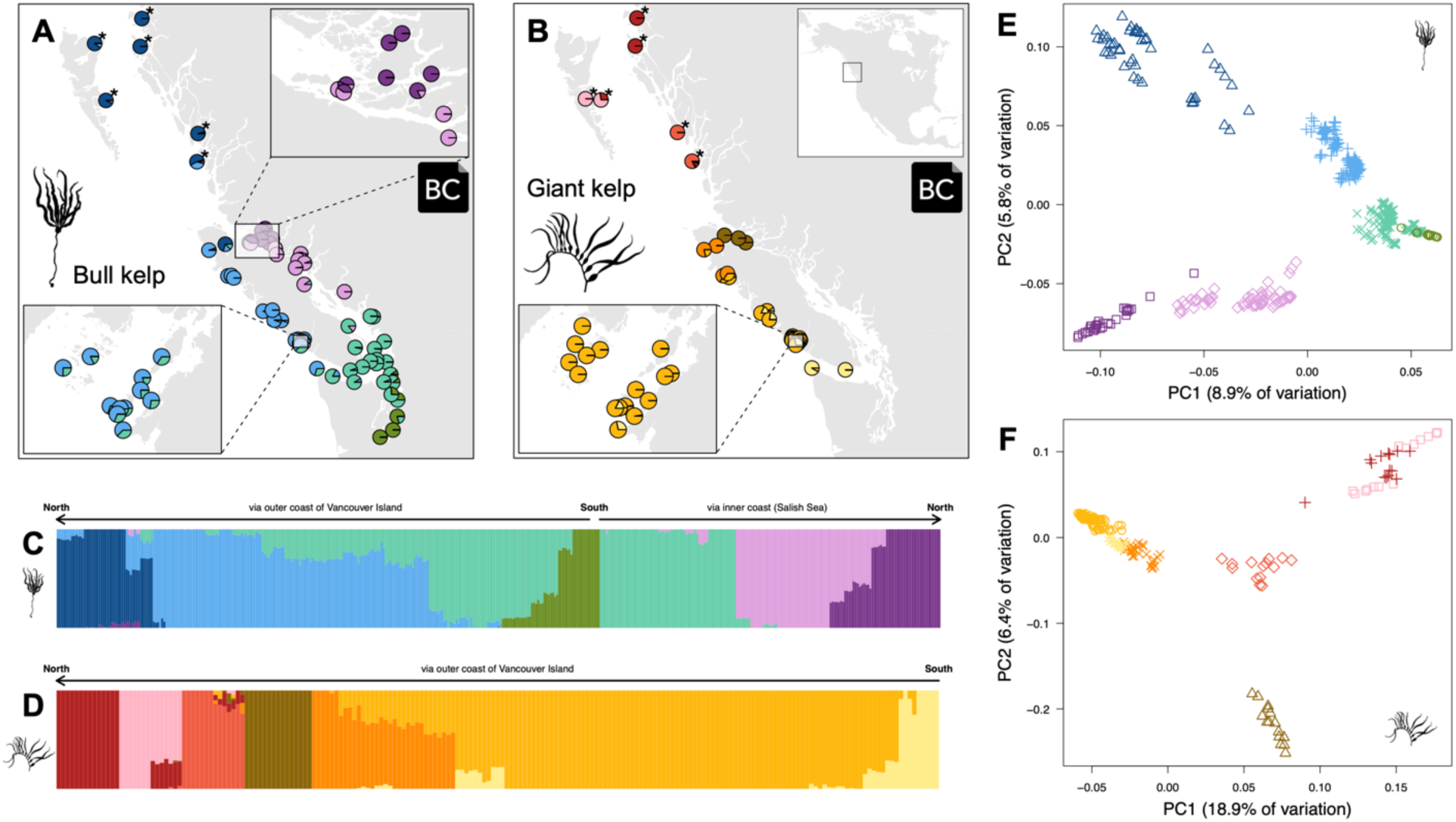
Genetic structure. Genetic structure of (A, C, E) bull kelp and (B, D, F) giant kelp in British Columbia and Washington. (A-B) Pie charts depict the proportion of ancestry in each population belonging to different genetic clusters, with clusters represented as different colours. An asterisk indicates locations that are approximate as coordinates were rounded to the nearest 0.5°. (C-D) The proportion of ancestry derived from each genetic cluster in each individual, with individuals represented as vertical lines. (E-F) Principal component analysis showing the clustering of individuals along the first two PC axes. Each point represents an individual, and is coloured according to the clusters from (A-D) and with a unique symbol for each cluster. Some samples and derived data are subject to a Biocultural (BC) Notice (see Resource Availability section and Table S1).

Some kelp populations in BCWA have remained stable in recent decades while others have experienced strong declines^2,4^. Small populations are often subject to multiple stressors that raise the risk of extirpation^27^. We assessed three genetic health indicators in each population: (1) effective population size (*N*_e_), with low *N*_e_ associated with higher extirpation risk due to demographic stochasticity, inbreeding, and drift^28^; (2) genetic diversity, which is required for populations to adapt to future challenges^29^; and (3) mean inbreeding coefficient, with higher inbreeding coefficients implying higher risks of inbreeding depression^30^. In bull kelp, *N*_e_ varied by more than two orders of magnitude (*N*_e_ = 33 to 6,236; Figure 2E,G), while nucleotide diversity (*π*) varied more than 40-fold between the highest-diversity populations (northern BC and northwest Vancouver Island) and the lowest-diversity population in southernmost Puget Sound (Squaxin Island, Washington) (Figure 2A). A similar geographic pattern was observed in inbreeding coefficients (*F*_ROH_100kbp_; Figure 2C). At least some of the variation in inbreeding coefficients is driven by high among-population variation in the rate of selfing (Figure S5).

**Figure 2.**
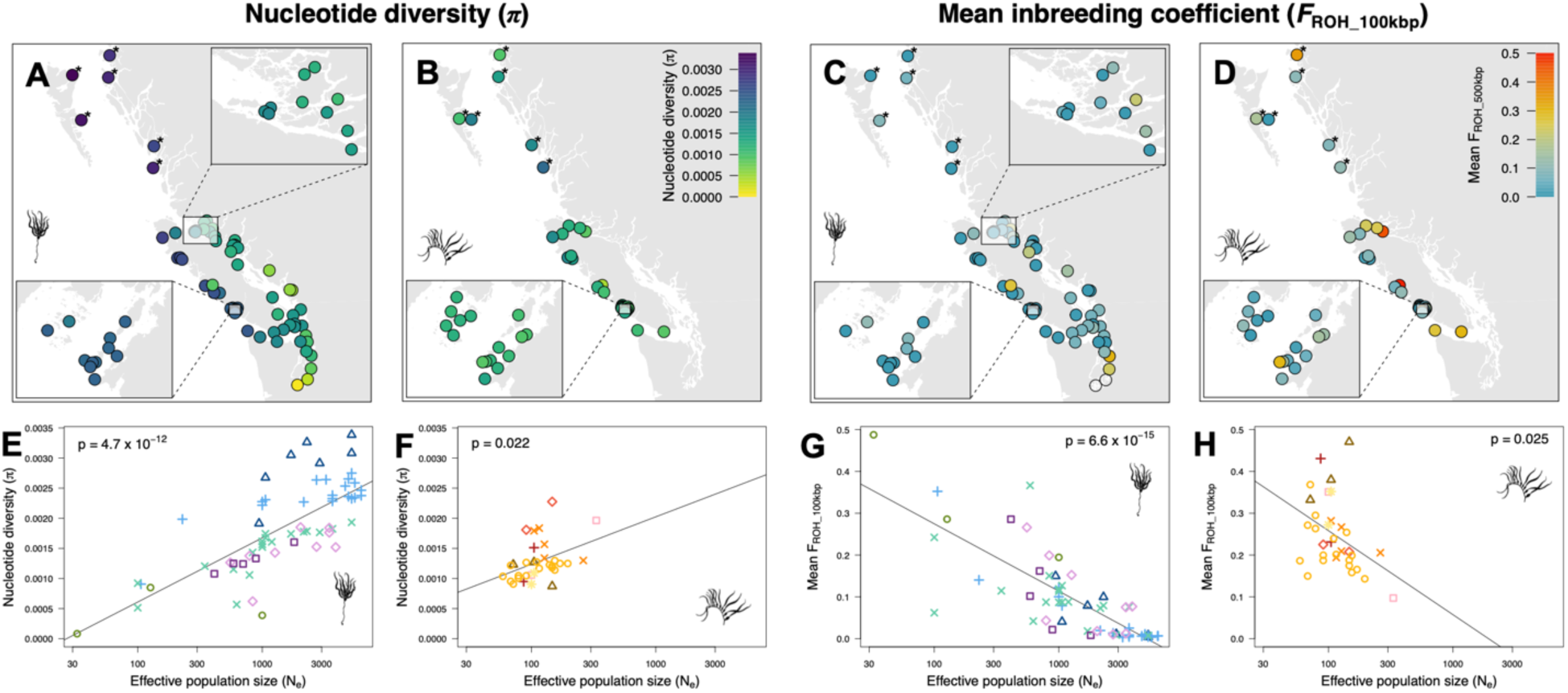
Genetic health indicators. Genetic health indicators for (A, C, E, G) bull kelp and (B, D, F, H) giant kelp. (A-B) Geographic distribution of nucleotide diversity (*π*). An asterisk indicates locations that are approximate as coordinates were rounded to the nearest 0.5°. (C-D) Geographic distribution of mean inbreeding coefficient (*F*_ROH_100kbp_). (E-F) Relationship between *π* and effective population size (*N*_e_). (G-H) Relationship between *F*_ROH_100kbp_ and *N*_e_. In (E-H), each point represents a population. Symbols and colours correspond to genetic clusters from Figure 1.

Selfing results when separate male and female haploid spores from a single diploid adult settle on the sea floor in close proximity (≤ 1 mm apart), a process facilitated by the limited dispersal distance of most spores^31–33^. In both species, the BCWA-wide selfing rate was 10%, in agreement with the ∼10% rate predicted for giant kelp from dispersal-based models^32^. In contrast to the extreme among-population variation observed in bull kelp, genetic health indicators were more uniform in giant kelp. Nucleotide diversity and *N*_e_ were comparatively low to moderate (Figure 2B,F) and *F*_ROH_100kbp_ was comparatively moderate to high in all populations (Figure 2D). As was the case for bull kelp, the highest-diversity populations tended to be in northern BC and northwest Vancouver Island (Figure 2B). In both species, *N*_e_ was positively correlated with *π* and negatively correlated with *F*_ROH_100kbp_ (Figure 2E-H). In addition, *N*_e_ was negatively correlated with selfing rate in bull kelp (Figure S6).

The frequent association of low effective population size with low genetic diversity and high inbreeding coefficients suggests that some populations face multiple genetic health risks that may lead to fitness declines and loss of evolutionary potential^27,34^. These populations tend to occur in Puget Sound, the Strait of Georgia and the inner Broughton Archipelago, as well as inner reaches of smaller water bodies away from the open coast (e.g., inner Quatsino Sound and Clayoquot Sound) (Figure 2). However, high inbreeding coefficients do not necessarily imply strongly reduced fitness, as natural selection may reduce inbreeding depression – narrowly defined here as a fitness reduction of inbred relative to outbred individuals from the same population^27^, though broader definitions are possible^35^ – by purging recessive deleterious alleles^8–10^. Specifically, in small populations where individuals are on average more closely related than in large populations^35^ (and where selfing is more common in bull kelp; Figure S6), alleles are more likely to be identical by descent, resulting in increased homozygosity of rare recessive deleterious alleles. This transformation of recessive deleterious alleles from a heterozygous state (‘masked load’) into a homozygous state (‘realized load’)^36^ exposes recessive deleterious alleles to natural selection. Over time, the loss of these alleles by purging can reduce the negative fitness consequences faced by inbred individuals^8^. However, inbreeding depression is unlikely to be entirely eliminated because natural selection is less effective in small populations^37^, making it difficult to purge alleles that are only mildly deleterious^8,38,39^.

Southern Californian populations of giant kelp experience substantial inbreeding depression^40–42^ but to our knowledge, inbreeding depression and purging have not been investigated in bull kelp nor in giant kelp from BCWA, where giant kelp forests are smaller^43^ and where *N*_e_ may sometimes be much lower (Figure 2) than in California (*N*_e_ = 50 to 2,500)^44^. We therefore looked for genomic signatures of purging by estimating genetic load in two ways. Firstly, we used Genomic Evolutionary Rate Profiling (GERP)^45^ to identify derived minor alleles (DMAs) at sites within the genome that are either evolutionarily labile or conserved across brown algae (Table S3), with DMAs at conserved sites considered more likely to be deleterious. Secondly, we annotated protein-coding genes using *SnpEff*^46^ to identify DMAs whose predicted impacts on proteins are either low, moderate, or high, and considered moderate- and high-impact sites to be more likely to be deleterious. We assumed that most putatively deleterious DMAs are likely to be at least partly recessive^47,48^.

We observed no evidence of purging in either species. We predicted that smaller populations would show a reduction in DMA frequency at evolutionarily conserved sites (GERP analysis) due to increased homozygosity and exposure to selection, yet DMA frequency was uncorrelated with population size (Figure 3A-B). Similarly, we predicted but failed to find evidence for a positive correlation between population size and frequency of moderate- to high-impact DMAs (*SnpEff* analysis; Figure 3C-D). Although purging has been empirically demonstrated in small populations of many species^10,49,50^, in many other cases it is not detected^51^, such as in recently bottlenecked populations where there has been insufficient time for purging to remove deleterious alleles^48^ or in extremely small populations where drift completely overwhelms natural selection^52^. Alternatively, if an ancestral population experienced a prolonged ancient bottleneck, most deleterious variants could have already been purged^53^. Such a bottleneck could conceivably have occurred during the Last Glacial Maximum, especially if recolonization of BCWA occurred from one or more small ice-free refugia, e.g., off the coast of Haida Gwaii as previously inferred for bull kelp^18^. Likewise, purging of alleles expressed during the haploid gametophyte life stage^54^ in all populations could reduce the signal of additional purging in diploid sporophytes that is brought about by small population size.

**Figure 3.**
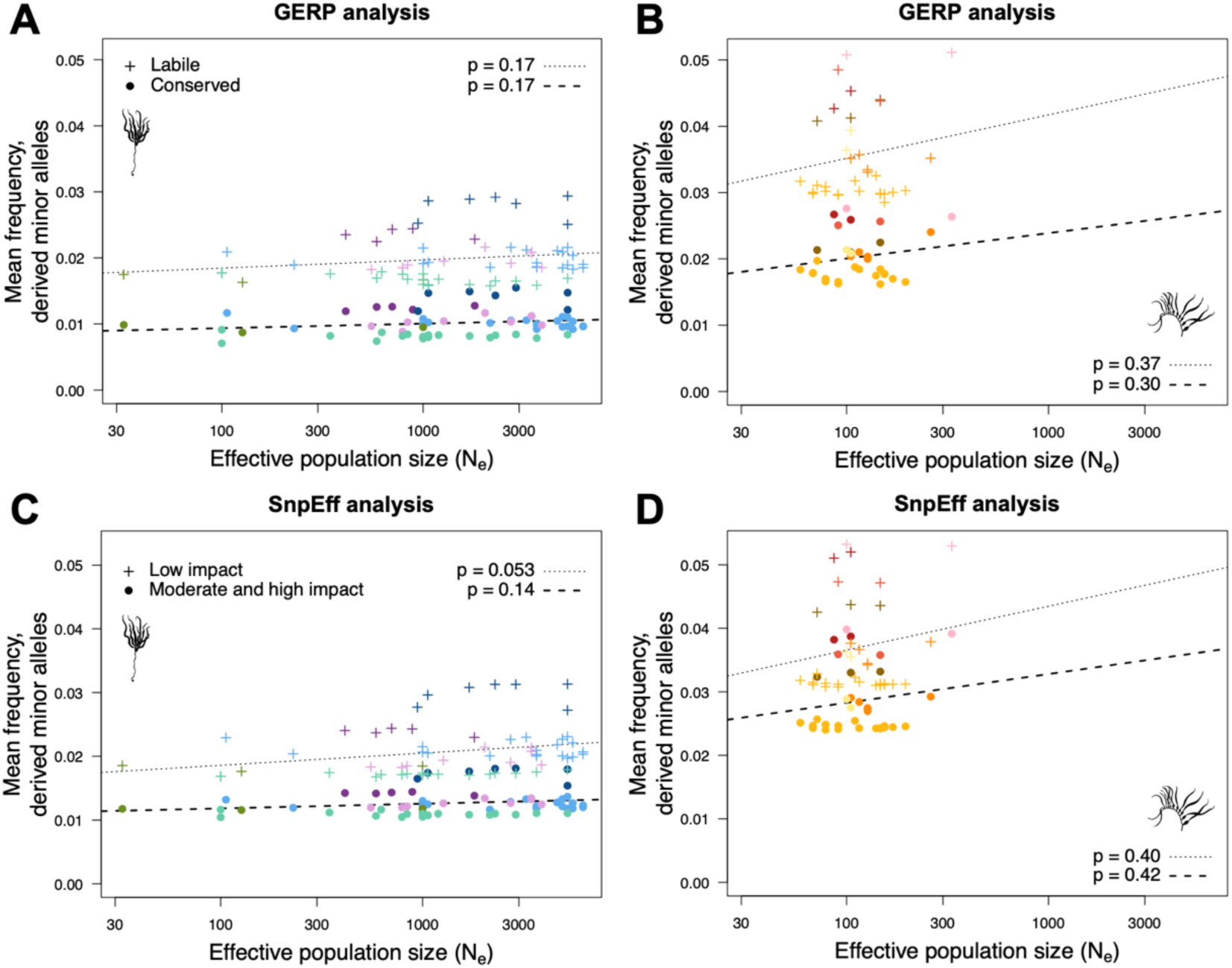
Lack of evidence for purging. Lack of evidence for purging by natural selection in (A-B) GERP and (C-D) *SnpEff* analyses. (A-B) Mean frequency of derived minor alleles (DMAs) does not vary with effective population size (*N*_e_) for either evolutionary labile (control) or conserved (putatively deleterious) sites in (A) bull kelp or (B) giant kelp. (C-D) Mean DMA frequency does not vary with *N*_e_ for sites predicted to have either a low (control) or moderate to high (presumed deleterious) impact on proteins in (C) bull kelp or (D) giant kelp. Each point represents a population. Symbols and colours correspond to genetic clusters from Figure 1.

Although we did not find evidence that natural selection is reducing inbreeding depression through purging, genetic drift may also reduce inbreeding depression within small populations^5,7,35,55^. Though rarely empirically tested (reviewed in ^7^; see also ^5,56^), this counterintuitive prediction’s theoretical foundations date back to Kimura *et al*.^37^, who noted that the more pronounced effects of genetic drift^57^ and reduced efficacy of natural selection in small populations would cause many mildly recessive deleterious alleles to become lost or fixed. With few recessive deleterious alleles segregating within a population, the relative difference in realized load between inbred and outbred individuals is expected to be very small^5,7,35,55^. Our results strongly support the expectations of this scenario in bull kelp. Firstly, the number of genomic sites with at least one copy of a DMA present in the population was reduced in small populations (Figures S7A, S8A), consistent with the expected loss of many DMAs. Loss of DMAs could occur through either drift or purging, yet the predictions of purging were not supported (see above). Secondly, remaining DMAs were present at higher frequency (Figures S7C, S8C) and more likely to be fixed (Figures S7E, S8E) in small populations. Thirdly, although the realized load of putatively deleterious DMAs was higher in selfed than outbred individuals in all populations (Figures 4A, S9A), the relative increase in realized load in selfed individuals was strongly positively correlated with population size (Figures 4B, S9B). These trends suggest that genetic drift has greatly reduced the relative fitness penalty for inbreeding relative to outcrossing within small populations. In contrast to the strong support for these predictions in bull kelp, trends were typically not significant for giant kelp (Figures 4, S7-S9), although lack of significance could relate to the minimal variation observed in population size.

**Figure 4.**
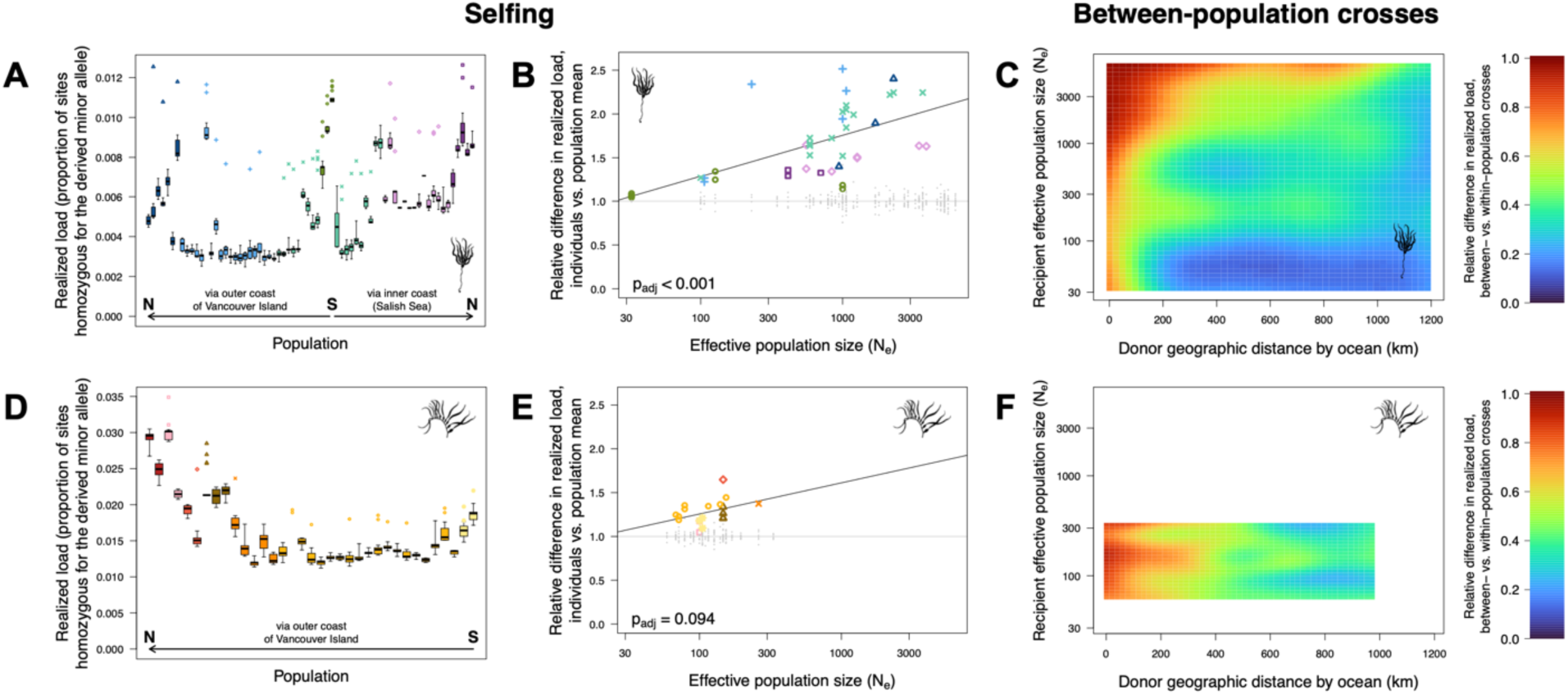
Predicted genetic load under different cross types, *SnpEff* analysis. Predicted changes in realized genetic load under different cross types for (A-C) bull kelp and (D-F) giant kelp suggest (A-B, D-E) a penalty for selfing and (C, F) reduced load in between-population crosses. Realized genetic load is measured as the proportion of sites homozygous for the derived allele at moderate- and high-impact sites in the *SnpEff* analysis. (A, D) Realized load is higher in selfed individuals (point symbols, one point per individual) than in non-selfed individuals (distribution in boxplots, with boxplot whiskers extending to the most extreme values). (B, E) Relative difference in realized load of selfed individuals (coloured symbols, one point per individual) compared to the mean realized load of non-selfed individuals, as a function of effective population size (*N*_e_). Non-selfed individuals are shown in small grey points for comparison. Adjusted p-values are calculated from 1,000 randomizations of *N*_e_. Symbols and colours correspond to genetic clusters from Figure 1. (C, F) Realized genetic load is predicted to be equal or lower (≤1.0) in between-population crosses (recipient x donor population) relative to within-population crosses (recipient x recipient). The predicted relative load (colour scale) is plotted as a function of recipient *N*_e_ and geographic distance by ocean to the donor population. The coloured two-dimensional surface was interpolated by kriging raw data points (Figure S12).

In addition to reducing inbreeding depression in small populations, the fixation of some recessive deleterious alleles due to genetic drift is expected to result in a fitness reduction known as ‘drift load’^5–7^. A large drift load could cause all individuals in small populations to exhibit reduced fitness relative to individuals from large populations^5,7,35,55^. We found strong support for increased drift load in small populations of bull kelp, as effective population size was positively correlated with the realized load of putatively deleterious DMAs in non-selfed individuals (Figure S10A,C). The relationship was not significant in giant kelp (Figure S10B,D).

The accumulation of drift load in small populations may also result in heterosis (i.e., hybrid vigour, or increased performance of between-population crosses relative to crosses within parental populations) due to the increased heterozygosity of recessive deleterious alleles expected in crosses between small populations that have independently and randomly fixed a different set of alleles^35,58^. Increased heterosis in small populations relative to large populations has been empirically demonstrated in numerous species^7,35,56,59,60^, though not universally^61^. Likewise, heterosis has been observed in some but not all Chilean populations of giant kelp, but an effect of population size was not tested^62^. Consistent with the expectations of heterosis, we frequently observed a lower realized genetic load in simulated between-population crosses (recipient x donor population) relative to simulated within-population crosses (recipient x recipient population) of both species (Figures 4C,F, S9C,F, S11, S12). As predicted, the realized genetic load was more strongly reduced in crosses in which the recipient population was smaller (bull kelp only; Table S4). In addition, realized genetic load was more strongly reduced when the donor population was more geographically distant (both species; Table S4), likely because geographically distant populations are expected to share few high-frequency deleterious alleles and so F_1_ homozygosity will be minimal.

Collectively, our analyses suggest a lack of purging in small populations of either species (Figure 3). Instead, genetic drift has caused the fixation of many putatively deleterious alleles (Figures S7, S8) in small populations of bull kelp, increasing the drift load of all individuals (Figure S10), reducing the difference in realized load between selfed and non-selfed individuals (Figures 4, S9) and leading to the potential for heterosis in between-population crosses (Figures 4, S9, S11, S12). Our findings result in several interesting evolutionary predictions that also have important implications for kelp conservation. These predictions are derived from genomic signatures and are based on the standard assumption that realized load is a reasonable proxy for fitness^8^, but we caution that we have not measured fitness of any individuals and experimental validation will be needed. Nonetheless, we predict that:

(1) All else being equal, individuals from small populations may perform more poorly in restoration or aquaculture than individuals from large populations of bull kelp. This prediction arises from the increased drift load observed in small populations (Figure S10) due to fixation of recessive deleterious alleles (Figure S7, S8) and presents an additional reason that large populations may be preferred as a genetic source over small populations above and beyond the higher genetic diversity of large populations (Figure 2E-F). However, a small population could still be a preferred genetic source if it is locally adapted to a historical environment that is a better match to the current or future environment of an outplanting site^63,64^.
(2) Offspring of crosses between small populations may perform better than offspring of crosses within a single small population. Any potential benefits of this predicted heterosis would need to outweigh the risks of outbreeding depression^65^ (e.g., through disruption of coadapted gene complexes^66^ or local adaptation^67,68^). The risks of outbreeding are poorly empirically known^65^ and often delayed until the F_2_ generation^65,66^, suggesting the need for long-term experimental monitoring. Nonetheless, the risk of outbreeding depression is generally minimal when populations occupy similar environments and have been recently fragmented (<500 years)^69^.
(3) Selfing may be an effective means of reproduction in small populations of bull kelp. In contrast, the fitness penalty for selfing is likely to remain high in large populations where selfing substantially increases the realized load relative to outbred individuals (Figures 4B, S9B). This situation could imply a shift from an effectively primarily outcrossing system to a mixed mating system as populations decrease in size and accumulate drift load (Figure S10), such as has been observed in leading-edge plant populations following range expansion^59^. Importantly, this shift could occur without the need to hypothesize explicit benefits to selfing such as reproductive assurance^70^ or the perpetuation of genotypes suited to local environments^71^. Due to the predicted fitness penalty for selfing in large populations, actions should be taken to minimize selfing (e.g., increasing the number of parents) in restoration cultures sourced from a large population. While outcrossing is still preferred when parents are sourced from a small population, avoiding selfing may be of lower priority, especially if obtaining high-quality reproductive material from numerous parents is difficult.

In summation, we have demonstrated strong genetic and geographic clustering in bull and giant kelp from BCWA that could aid in designating MUs or seed-transfer zones. Small populations face multiple genetic health risks but show no evidence of purging. Instead, allele-frequency changes in small populations appear to be dominated by genetic drift. Our genomic analyses have revealed fundamental insights into the evolutionary dynamics of small populations and imply several strategies that could be cautiously applied (pending experimental validation) to conservation and restoration of these at-risk and declining^1–4^ kelp species.

## Methods

### EXPERIMENTAL MODEL AND SUBJECT DETAILS

This study generated sequencing data for 449 bull kelp and 239 giant kelp. Sample sites and sample sizes are provided in Table S1. Each sample site corresponds to a single kelp forest (hereafter, “population”), sampled between 2021 and 2023. For up to 15 individuals from each population, a cutting of blade tissue was collected and dried using silica gel. Effort was made to collect from blades a minimum of 3 m apart, as assessed from the water surface, but the spacing of holdfasts along the ocean floor could not typically be determined for either species and 3 m spacing could not be ensured for some of the physically smallest bull kelp populations.

### METHOD DETAILS

#### DNA extractions and sequencing

We extracted DNA from dried blade tissue using a cetyltrimethylammonium bromide (CTAB) protocol, substantially modified from one developed by ^73^. We disrupted approximately 0.5 cm^2^ of tissue using a *Qiagen* TissueLyser, then added 1 mL of citrate wash buffer (0.055 M sodium citrate dihydrate, 0.030 M EDTA, 0.150 M NaCl, adjusted to pH 8.0)^74^ and incubated with agitation for 15 min at room temperature. We then centrifuged at 10,000 G for 5 min, discarded the supernatant, and repeated the wash step a second time but without the 15-min incubation. To the resulting precipitate, we added 700 uL of CTAB isolation buffer (1% CTAB, 5 M NaCl, 0.5 M ethylenediaminetetraacetic acid [EDTA], 1% polyvinylpyrrolidone [PVP], 10 mM Tris-HCl), 14 uL proteinase-K (20 mg/mL) and 10 uL of RNAse A (100 mg/mL), and incubated at 55°C, inverting the sample tube every 5-10 min for the first ∼40 min before leaving it to incubate overnight. The next day, we centrifuged for 10 min at 13,300 G, resulting in the formation of an aqueous clear upper layer and a green lower layer. We transferred the upper layer (∼650 uL) to a new tube and added 0.2-0.3 volumes of 100% ethanol very slowly with constant stirring to precipitate residual polysaccharides without precipitating DNA. To this solution, we then added 650 uL of 24:1 chloroform:isoamyl alcohol, mixed briefly by inversion, and centrifuged at 12,000 G for 5 min. We then transferred 600 uL of the resulting upper layer to a new tube (leaving ∼50 uL behind to prevent contamination of the transferred portion with the bottom layer), added 600 uL of isopropanol to the transferred upper layer, vortexed the solution, and centrifuged at 13,300 G for 30 min at room temperature. After removing the supernatant, a small white DNA pellet remained, which we washed twice with 250 uL of 70% ethanol. We air-dried the pellet and dissolved it in 50 uL of nuclease-free water. We then cleaned the DNA using magnetic beads (*Sergi Lab Supplies*) at 0.8X following the manufacturer’s protocol.

Initial testing indicated that the functionality of cleaned extractions in downstream applications could not be reliably predicted from standard quality checks (i.e., using a *Thermo Scientific* NanoDrop spectrophotometer and a *Thermo Scientific* Qubit fluorometer), suggesting undetected contaminants in some samples. We therefore ran a test polymerase chain reaction (PCR) on each sample to rule out the presence of PCR inhibitors. We used primers for the internal transcribed spacer (ITS) locus for each species (bull kelp: *N. luetkeana* F and *N. luetkeana* R2 from ^75^; giant kelp: KG4 reverse and *Macrocystis integrifolia* forward primers from^76^). PCRs were performed in a 5-uL reaction volume using 2.5 uL of *Roche* 2X KAPA Hifi HotStart ReadyMix, 0.15 uL of 10 uM forward primer, 0.15 uL of 10 uM reverse primer, and 1.7 uL of nuclease-free water. Reactions were denatured at 95°C for 3 min; followed by 35 cycles of 98°C for 20s, 62°C for 15s and 72°C for 30s; and a final extension of 72°C for 2 min. We visualized samples on a 2% agarose gel to check for the presence of a 575-bp band in bull kelp^75^ and a 912-bp band in giant kelp^76^. Samples without a strong band were re-extracted or a different individual from the same population was selected instead.

We prepared libraries for whole-genome sequencing (WGS) using *Integrated DNA Technologies (IDT)* xGen DNA Library Prep EZ Kits, following the manufacturer’s protocol with the following options and minor modifications: 165 ng of input DNA topped up to 16 uL in nuclease-free water; use of the *IDT* xGen Deceleration Module to slow enzymatic fragmentation, with reaction times reduced by 1 min relative to the timing indicated for each specific lot of each *IDT* kit (typically 12 min reduced from an indicated 13 min); the use of *IDT* xGen 10-nucleotide primers; no use of the optional *IDT* xGen Normalase Module; and six PCR cycles to account for the additional cycling requirements of the Deceleration Module. We performed bead-cleaning steps using *Sergi Lab Supplies* magnetic beads at the same ratios indicated in the protocol for AMPure XP beads (*Beckman Coulter*). We combined between 62 and 66 barcoded samples per library pool (except for initial test libraries of fewer samples) across a total of 12 library pools. We sent samples to Génome Québec (Montreal) or Canada’s Michael Smith Genome Sciences Centre (Vancouver) for 150-bp paired-end sequencing on an *Illumina* NovaSeq 6000, targeting 2.5 billion paired-end reads per library pool.

Surprisingly, many of our library pools failed quality control at the sequencing core due to the presence of a large, unexpected secondary peak of short fragments (ca. 310-330 bp; target peak size typically >700 bp) that was not detectable in gel electrophoresis on a manual agarose gel but appeared on the results of automated electrophoresis machines at the sequencing core. This secondary peak was repeatably detected across multiple independent sample submissions, was present in multiple library pools prepared months apart, could not be removed by further bead clean-up at stringent bead:sample ratios, and could not be visualized in house on any manual agarose gel under any visualization settings, suggesting that it may have represented an unknown contaminant rather than short DNA fragments. Test sequencing of affected libraries on an *Illumina* MiSeq revealed no apparent negative impacts on the sequencing reaction and a size distribution of sequenced fragments corresponding to the primary (target) peak only. We therefore proceeded with sequencing all of our library pools and could detect no impacts on the quality or quantity of the final data we obtained.

#### Alignment to reference genome

Prior to aligning raw reads to reference genomes, we trimmed adapter sequences, merged overlapping paired-end reads, and performed basic quality and read-length filtering using *fastp* v.0.23.2^77^ with default parameters. We added a unique read-group to each sample and aligned filtered, adapter-trimmed reads to reference genomes for either bull kelp (467.6 Mb; NCBI assembly GCA_031213475.1)^78^ or giant kelp (537.5 Mb; JGI PhycoCosm assembly *Macrocystis pyrifera* CI_03 v1.0)^79^ using *bwa-mem* v.0.7.17-r1188^80^ with default parameters. We then sorted and merged aligned reads (paired, unpaired, and merged) into a single file for each sample using SAMtools v.1.17^81^, and removed read duplicates using *Picard* v.2.26.3^82^. We removed one giant kelp individual that had an extremely low read alignment rate (1.4%) and for which the tissue sample was noted to have been visibly degraded and contaminated with symbionts.

To correct potential misalignments around indels that could lead to the identification of false variants, we realigned reads using *GATK* v.3.8^83^. We first identified target intervals representing putative indels using the *RealignerTargetCreator* tool from *GATK*, identifying targets from all individuals in a single command for each species. We then used the *IndelRealigner* tool from *GATK* to realign reads around indels.

#### SNP genotyping

We performed SNP genotyping steps separately for datasets of bull kelp from BCWA only, giant kelp from BCWA only, and giant kelp from all global samples. We identified Single Nucleotide Polymorphisms (SNPs) and called genotypes using *BCFtools* v.1.11-1.19^81^. We created binary Variant Call Format (VCF) files of genotype likelihoods using *bcftools mpileup*, requiring minimum base and mapping qualities of 30 (*-Q 30 -q 30*), and called genotypes with *bcftools call* using the multiallelic caller (*-m*) and outputting variant sites only (*-v*). We filtered raw genotype calls with *bcftools view* to include only SNPs (*-v snps*) that were biallelic (*-m 2 -M 2*) and with both alleles observed in our samples and not only in the reference genome (*-q 0.000001:minor*). We retained only nuclear SNPs by removing the (*-t ^)* mitochondrial (mtDNA) and chloroplast (cpDNA) assemblies, as identified in the reference genomes.

We next filtered raw SNP calls to generate a set of high-quality sites for downstream analyses. We required a minimum depth of 10 to retain a SNP call by using the *+setGT* plugin of *BCFTools* to set genotypes to missing at sites where depth was less than 10. We then used *bcftools view* to exclude sites with heterozygosity above 75%. We also removed sites that failed any of five additional quality tests flagged by *BCFTools* in the INFO column of the VCF files. Specifically, we retained only sites where the following was true: log-likelihood ratio of Segregation Based Metric (SGB) ≥ 2, p-value of Mapping Quality Bias (MQB) ≥ 0.05, p-value of Mapping Quality vs Strand Bias (MQSB) ≥ 0.05, p-value of Read Position Bias (RPB) ≥ 0.05, and p-value of Variant Distance Bias (VDB) ≥ 0.05. These quality metrics and their cutoff values were selected based on inspecting heterozygous SNP calls in selected regions identified as putative Runs of Homozyosity (ROHs) from visual inspection of sliding windows of heterozygosity in unfiltered SNP datasets (i.e., from inspecting presumed erroneous SNPs). We additionally excluded sites above the 98^th^ percentile of depth of coverage across all samples combined.

After applying SNP filters, we removed seven bull kelp and nine giant kelp individuals from downstream analyses due to high missing data (≥ 50% missing). The mean depth of coverage of all retained samples was ≥ 9.5X. We then recalculated allele frequencies (*-t AN,AC*) using the *+fill-tags* plugin of *BCFTools*. For giant kelp we recalculated allele frequencies and created datasets for separate subsets of individuals (rangewide, North American, and BCWA samples, respectively).

We then used *BCFTools* to perform additional site filtering, retaining sites with a minimum minor allele frequency of 0.01 (-q 0.01:minor) and maximum 20% missing data. We also retained only putative autosomal SNPs by removing sites on putative sex chromosomes (JARUPZ010000001.1 in *Nereocystis* and scaffold_2 in *Macrocystis*; sex chromosomes were identified as scaffolds were large regions contained approximately half the depth of coverage of the remainder of the genome across all individuals, suggestive of a haploid region). To retain only a high-quality nearly scaffold-level assembly, we removed scaffolds and contigs smaller than 1.5 Mbp (cutoff selected by visual inspection of scaffold lengths), resulting in retention of 94% and 87% of the putatively autosomal *Macrocystis* and *Nereocystis* genomes, respectively. Because some downstream applications required the removal of closely-related individuals, we inferred relatedness between all pairs of individuals using *ngsRelate* v.2^84^. We subsetted our autosomal dataset to a minimum minor allele frequency (MAF) of 0.05 (*bcftools view -q 0.05:minor*) and thinned to a minimum distance of 10 kbp between SNPs using *VCFtools* v.0.1.16^85^and used the resulting VCF file to run *ngsRelate* with default parameters, using genotype likelihoods rather than called genotypes. We used the KING-robust kinship estimator (calculated from the 2D site-frequency spectrum^86^) to identify individuals that were genetically identical or first-degree relatives (i.e., parent-offspring and full-sib pairs), with thresholds for delimiting relatedness categories following ^87^. We then selected the minimum set of individuals that needed to be excluded to create two sets of samples with no genetically identical individuals and additionally no first-degree relatives, respectively, retaining the individual with the higher depth of coverage from pairs where the choice of individual to exclude was arbitrary. From the un-thinned files with minimum MAF of 0.01, we created multiple datasets excluding 13 genetically identical and three additional first-degree relatives for bull kelp respectively, and 23 genetically identical and 16 additional first-degree relatives for giant kelp, respectively. As expected, all pairs of close relatives were from the same sampling site. Genetically identical individuals may reflect asexual reproduction, which has been reported in giant kelp^43,88^, or errors in sampling in which the same individual was sampled more than once (despite a minimum distance implemented between individuals) because it was sometimes difficult to tell from the surface of the water which blades belonged to which individual.

Finally, after removing close relatives, we thinned SNPs to a minimum distance of 10 kbp using *VCFtools*. We confirmed that 10 kbp was an appropriate thinning distance based on visual inspection of linkage-disequilibrium (LD) decay curves showing that LD had substantially declined from its maximum and 10 kbp was near the inflection point in the curve. We calculated LD using a 2% random sample of pairwise comparisons (*--rnd_sample 0.02*) using *ngsLD* v.1.2.1^89^, and plotted LD decay curves (with LD measured as r^2^) using the *fit_LDdecay.R* script from *ngsLD*.

The above methods describe the biallelic SNP dataset(s) used in the majority of analyses, yet some analyses required information about all sites in the genome. We therefore also called genotypes at invariant and triallelic (or quadrallelic) sites using the same procedure as for our biallelic SNPs with minor modifications. Specifically, we repeated our initial *bcftools call* command with the variant flag (*-v*) removed to call genotypes at all sites in the genome. We identified invariant sites as those with only one allele observed in our samples (*bcftools view -Q 0.000001:nonmajor)* and excluded indels (*-V indels*). Subsequent filtering steps were the same as for biallelic SNPs except that the heterozygosity filter and variant quality filters (SGB, MQB, MQSB, RPB, and VDB) were not applied, nor was filtering for a minimum MAF. We identified triallelic (or quadrallelic) sites as SNP variants with a minimum of 3 alleles (*bcftools view -m 3 -v snps*), and implemented all filters except for a minimum MAF. Finally, we also reprocessed our biallelic SNPs in the same way as previously described, except without any minimum MAF, to generate a set of all biallelic SNPs in the genome regardless of frequency. We then used *bcftools concat* to combine the separately-filtered invariant, biallelic, and triallelic (or quadrallelic) sites into a single dataset representing all high-quality filtered sites in the genome.

#### Derived minor allele classification

We identified and classified derived minor alleles (DMAs) into different categories of deleteriousness using (1) Genomic Evolutionary Rate Profiling (GERP)^45^ and (2) the genetic variant annotator *SnpEff*^46^.

(1) Our GERP analysis closely followed ^48^. GERP consists of inferring the rates of evolution at individual sites in the genome across a phylogeny of closely related species, to identify sites that are relatively conserved across taxa (where new mutations are more likely to be deleterious) and sites that evolve relatively rapidly (where new mutations are less likely to be deleterious)^45^. We obtained a time-calibrated composite phylogeny of brown algae (Phaeophyceae) using ^90^ as a backbone, primarily for deep divergences within the class (e.g., between orders). We added additional branches and divergence times to the backbone using phylogenies that focused on subsets of brown algae^91–95^. For kelp specifically, we used the divergence times and branching structure from ^95^. All other branch times added to the composite brown algae phylogeny were also as given in original publications, except that divergences between *Choristocarpus* and *Discosporangium* and between *Halopteris* and *Sphacelaria* from ^94^ were proportionally rescaled relative to the earliest split within Phaeophyceae (Discosporangeales vs. others) according to ^90^, to account for the substantially older early divergence times in ^90^ relative to ^94^.

We downloaded reference genomes for species in the brown algae phylogeny (accession numbers in Table S3) and split the genomes into shorter fragments of 500 bp using the *reformat.sh* script of *BBmap* v.39.06^96^. We then aligned fragments to the focal species’ (bull kelp or giant kelp) genome following ^48^ using *bwa-mem* v.0.7.17-r1188^80^ with modified mismatch penalty (*-B 3*) and gap opening penalty (*-O 4,4*); used the *view* command in SAMtools v.1.17^81^ to filter by mapping quality (*-q 2*) and remove reads aligning to multiple locations (*-F 2048*); and converted alignments to fasta format using the *pileup* command in *HTSBox* v.r345^97^, requiring minimum mapping and base qualities (-q 30 -Q 30), length (-l 35) and depth (-s 1), and printing a random allele (-R). If multiple alignments were available that mapped to the same branch on the phylogeny (e.g., multiple species in the same genus), the alignment with the highest number of primary mapped reads was retained and other alignments discarded. We then used a custom script heavily modified from ^48^ to combine all species into a single alignment for each scaffold or contig.

To calculate GERP scores for each site (positive scores indicate fewer substitutions than expected across the phylogeny, i.e., higher evolutionary constraint), we used the modified *gerpcol* script (v.2023/11/20) from ^48^ to run *GERP++*^98^. We used our previously-calculated (see above) species-specific transition:transversion ratios (*-r*) and a brown algae substitution rate of 8.135 × 10^−4^ mutations per million years (i.e., per unit of branch length; *-s 0.0008135*). This substitution rate was derived by averaging substitution rates for *Ectocarpus* and *Scytosiphon* (4.07 × 10^−10^ and 1.22 × 10^−9^ substitutions per generation, respectively^99^) and assuming a generation time of one year. We also printed the alleles for each site (*-e*) at the three closest outgroups (either bull or giant kelp, *Saccharina japonica* and *Laminaria digitata*) for each species. Finally, we used the GERP scores (S) to classify sites as evolutionarily conserved (S > 0.5) or evolutionarily labile (S ≤ 0.5), retaining only sites with sufficient species alignments to expect a more than 0.5 substitutions (N > 0.5) across the entire phylogeny.

To incorporate the GERP scores with our SNP datasets, we used custom *R* scripts to determine the ancestral and derived alleles at each SNP site, and calculate the derived allele frequency globally and in each population. Using the alleles printed for the three closest outgroups, we considered an allele to be derived if it was not present in any of the outgroups (allowing for up to two outgroups to have missing data). If neither or both alleles were present in the three outgroups, ancestral and derived alleles could not be defined. Because genetic load is only relevant to putatively deleterious mutations, we excluded sites with a global derived allele frequency greater than 0.5 as these sites are unlikely to be deleterious. We obtained a total of 18,905 SNPs (9,396 labile and 9,509 conserved) for downstream analyses in bull kelp and 9,074 SNPs (4,884 labile and 4,190 conserved) in giant kelp.

For our second method of classifying sites, we used *SnpEff* v.5.2a^46^ to predict the impacts of derived mutations in protein-coding genes. *SnpEff* classifies predicted protein impacts as low, moderate, or high, with an additional modifier category for non-coding variants^46^. Gene annotations were available in the reference genome for giant kelp but not bull kelp. We transferred gene annotations from giant kelp to bull kelp using *Liftoff* v.1.6.3^100^ using the options *-infer_genes -copies -a 0.95 -s 0.95 -d 5.0 -flank 0.8 -polish*, and retained only annotations flagged with a valid open reading frame. For both species, we built *SnpEff* databases from gene annotation files (*-gff3*) using the *build* command, and then added variant annotation to VCF files of SNP datasets with only identical individuals removed using the *ann* command with default parameters. We used custom scripts to calculate global and population-specific ancestral and derived allele frequencies of annotated variants, using the same ancestral and derived definitions as determined above in the GERP analysis. We obtained a total of 115,549 non-modifier SNPs (51,988 with low impact and 63,561 with moderate or high impact) for downstream analysis in bull kelp and 254,297 SNPs (109,642 with low impact and 144,655 with moderate or high impact) in giant kelp.

### QUANTIFICATION AND STATISTICAL ANALYSIS

#### Genetic structure

We performed principal component analysis using *SNPRelate* v.1.38.0^101,102^ with default parameters, using the SNP datasets with up to first-degree relatives removed. We also performed genetic clustering analyses with *fastSTRUCTURE* v.1.0^22^, using the SNP datasets with up to first-degree relatives removed. We used simple priors and varied the number of clusters (K) from two to 10. For each K-value, we ran the program 100 times and selected the run with the highest likelihood. We used these 10 highest-likelihood runs to select the optimal range of K-values using the *chooseK.py* script distributed with *fastSTRUCTURE*. For bull kelp, the optimal value was K=8 by both reported criteria, but two of the clusters represented only one or two low-diversity populations in a small, isolated geographic area. As these two clusters may have represented differentiation due to recent bottlenecks rather than long-term regional genetic structure, we opted to use K=6 as the best model to represent regional genetic structure across BCWA. For giant kelp from BCWA only, the optimal value was K=7; from North America only, optimal K=7; from all global samples, optimal K was between 5 and 6 but we selected K=5 because K=6 contained one virtually unused cluster for which no individual had >1% ancestry.

We used the above-identified clusters to group populations for calculating pairwise genetic differentiation and divergence within BCWA for each species, and between the Southern and Northern Hemispheres in giant kelp (irrespective of ecomorph identity). In giant kelp, we also calculated statistics between BCWA (all presumably *integrifolia* ecomorph) and California *pyrifera* ecomorphs, excluding the one high-quality *integrifolia* ecomorph for which data were available from California (Figure S2). We calculated pairwise genetic differentiation (Weir and Cockerham’s^103^ *F*_ST_) using *hierfstat* v.0.5-11^104^, using the SNP datasets with up to first-degree relatives removed. We calculated pairwise genetic divergence (*d*_XY_, the average number of nucleotide differences between two random individuals from different populations^105^) using *pixy* v.1.2.7.beta1^106^, using the SNP datasets with only identical individuals removed and containing both variant and invariant sites.

To test for a pattern of isolation by distance, we plotted *d*_XY_ against the geographic distance between populations by the shortest ocean route (km). The geographic distance was calculated by converting a polygon of the BCWA coastline^107^ into a raster at 1-millidegree resolution using the *rasterize()* function in the *R* package *raster* v.3.6-26^108^. This high resolution was required to accurately represent connectivity along BC’s complex coastline, but calculations between very distant populations became computationally intractable. We therefore split the coastline into two regions: (1) the Salish Sea and Vancouver Island and (2) northern BC. In northern BC we coarsened the resolution to 2-millidegrees to aid computation. We manually inspected both rasters to ensure that population sampling locations were accurately rasterized as ocean and not land and that narrow passages between islands through which kelp might disperse were fully passable, manually converting pixels from land to ocean if needed. We used the *R* package *gdistance* v.1.6.4^109^ to calculate an 8-directional transition matrix for each raster using the *transition()* function; to correct the transition matrix, to account for the fact that degrees of latitude and longitude are not equal in distance, using *geoCorrection*() with type “c” correction; and to calculate the least-cost path between populations using *costDistance().* For population pairs where one population was located in each of the two rasters, we calculated the distance between each population and an intermediate coastal point shared between the two rasters (Cape Caution, BC; 51.165°N, 127.797°W) and then summed the two distances. This method provided computational tractability and also forced populations to disperse along the central coast of BC rather than across the open waters of Queen Charlotte Sound, which is likely a biologically realistic representation given the assumption of stepping-stone dispersal between populations. After obtaining geographic distances, we performed linear regressions of the relationship between *d*_XY_ and geographic distance using the *lm()* function in *R* v.4.2.3^110^. Unless stated otherwise, all simple linear regressions in this study were also performed with *lm()*. To obtain adjusted p-values for the relationship between d_XY_ and geographic distance, we performed Mantel tests^111^ by permuting the geographic distance matrix 1,000 times.

#### Genetic health indicators and selfing rate

We calculated three genetic health indicators for each population: (1) We estimated effective population size (*N*_e_) using *roh-selfing* v.2024/02/23^112^. Because *roh-selfing* requires information on runs of homozygosity (ROHs) as input, we masked repetitive regions of the genome to ensure that potential misidentified SNPs in these regions would not prevent accurate inference of ROHs. We identified repetitive regions in the reference genomes with *Red*^113,114^ using the *Red2Ensembl.py* script distributed with *Ensembl Plants* v.1.2^115^, and then masked them in the SNP datasets with only identical individuals removed using the *intersect* command in *BEDTools* v.2.30.0^116^. We then further masked potentially problematic regions of the genome by excluding short heterozygous regions surrounded by ROHs. To do so, we estimated an initial round of ROHs for each individual in each population from the repeat-masked SNP datasets using the *roh* command of *BCFTools* v.1.19^81^, with allele frequencies calculated automatically for populations of ≥4 individuals by *BCFTools*, or else using global allele frequencies across all individuals (provided with the *--AF-file* flag) for populations with <4 individuals. We then used a custom *R* script to identify short runs of heterozygosity (ROHets; the inverse of ROHs) ≤10 kbp in length and surrounded by ROHs in each individual. We also used a custom *R* script to identify 10-kbp windows of the genome that contained a short ROHet in greater than *n* individuals, where *n* was equal to the 99.99^th^ percentile of a Poisson distribution with parameter *λ* equal to the mean number of individuals containing a short ROHet across all 10-kbp windows of the genome. Finally, we masked these windows (in addition to the mask for repetitive regions) in our SNP datasets by re-running *Red* and *BEDTools* as described above. Using these masked SNP datasets, we then estimated a final round of ROHs for each individual in each population using the *roh* command of *BCFTools* as described above, retaining results only for populations of ≥4 individuals.

In addition to ROHs, *roh-selfing* requires estimates of inbreeding coefficient (*F*) and Tajima’s^117^ *D* as input. For each population, we calculated *F* using *PLINK* v.1.90b6.21^118^ and Tajima’s *D* using *VCFtools* v.0.1.16^85^. We then ran the *RF-sequential* model (model ID 202310021917048AtJy) of *roh-selfing*^112^ on each population. We generated a single estimate of *N*_e_ for each population by taking the mean of log_10_(*N*_e_) for each autosomal scaffold or contig.

(2) We calculated nucleotide diversity (*π*)^105^ for each population using *pixy* v.1.2.7.beta1^106^, using the SNP datasets with only identical individuals removed and containing both variant and invariant sites.

(3) We calculated mean inbreeding coefficients *(F_ROH_*) for each population, representing the proportion of the genome that is in ROHs. The estimation of ROHs using *BCFTools* was described above. We then calculated *F*_ROH_100kbp_ for each individual as the summed length of ROHs ≥100 kbp divided by the total length of all scaffolds and contigs for which ROHs were inferred, and calculated the mean across individuals within each population.

In addition to these three genetic health indices, we also calculated the observed selfing rate in each population from long ROHs (≥ 500 kbp), with selfed individuals expected to have approximately 50% of their autosomal genomes in long ROHs. We used *BCFTools* above to calculate ROHs for *roh-selfing* as the program is trained on *BCFTools* output, but were concerned that any false SNP regions not removed by our SNP filtering and masking procedures might break up long ROHs, making it difficult to infer accurate selfing rates. We therefore identified long ROHs using *ROHan* v.1.0.1^119^, which classifies ROH status in large windows, does not rely on called genotypes, and can accommodate a background heterozygosity rate in putative ROHs. For each individual, we ran *ROHan* using indel-realigned BAM files (described in the *Alignment to reference genome* section), restricting the analysis to large autosomal scaffolds (*--auto*), using 500-kbp windows (*--size 500000*), with an expected background heterozygosity rate in non-ROH regions (*--rohmu*) of 5.0 × 10^−4^ for bull kelp and 6.0 × 10^−4^ for giant kelp, and supplying a species-specific transition:transversion (TSTV) ratio (*--tstv*). The TSTV ratio was 1.45 for bull kelp and 1.75 for giant kelp, calculated using *bcftools stats* from the SNP datasets with up to first-degree relatives removed. The background heterozygosity rate was determined heuristically for each species by running *ROHan* on several individuals strongly suspected of being selfed (based on visual inspection of Manhattan plots of observed heterozygosity, described below) and plotting histograms of the heterozygosity inferred by *ROHan* across all 500-kbp windows (with at least 80% of the sites having data). For selfed individuals, the distribution of heterozygosity is expected to be bimodal, with two large peaks corresponding to windows in ROHs and not in ROHs, respectively. The background heterozygosity rate was selected to approximately correspond to the maximum value of the first peak of this distribution. After running *ROHan* with optimized parameters for each individual, the mean *F*_ROH_500kbp_ was calculated for each individual as the proportion of 500-kbp windows inferred to be in ROHs using *ROHan*’s mid-value estimates of heterozygosity.

Selfing rate was then calculated as the proportion of individuals in each population (using non-identical individuals only) with *F*_ROH_500kbp_ > 0.3536. The expected *F*_ROH_ of selfed individuals is 0.5, while parent-offspring pairs and full siblings are expected to have *F*_ROH_ = 0.25. We used 0.3536 as the threshold for binary classification following a proposed inference criterion for distinguishing 0.5 and 0.25 kinship coefficients^87^. In two bull kelp populations (NL-PS-06 and NL-PS-07), selfing rate could not be reliably determined from *F*_ROH_500kbp_ because *ROHan* was unable to classify most segments of the genome as either ROH or non-ROH given that the entire genome had extremely low genetic diversity. For these populations, we additionally used the criterion *F*_ROH_100kbp_ > 0.3536 from *BCFTools* to identify individuals that could potentially be selfed. We then confirmed the inference of selfing by visual inspection of Manhattan plots of observed heterozygosity (*H*_o_) in 100-kbp windows across the genome, calculated using *pixy* as described above for calculating *π*. We visually confirmed the expected presence of ROHs spanning entire chromosomes or the majority of chromosomes in selfed individuals. In addition, in one giant kelp population (MP-NC-02), all individuals had high *F*_ROH_500kbp_, suggesting that non-selfed individuals could potentially exceed the 0.3536 threshold. We reclassified four individuals from this population as non-selfed after visual inspection of Manhattan plots suggested that the genomes had numerous smaller ROHs, but few ROHs approaching chromosome length.

#### Tests for purging and genetic drift

For both GERP and *SnpEff* analyses, we used custom scripts to test for purging and examine the effects of genetic drift in small populations. We expected that purging would remove putatively deleterious alleles from small populations but have no effect on frequencies of alleles in less deleterious categories. We considered derived minor alleles (DMAs) at evolutionarily conserved sites to be putatively deleterious in GERP analyses and DMAs at moderate- and high-impact sites to be putatively deleterious in *SnpEff* analyses. We calculated the mean frequency of DMAs in each allele category from sites genotyped at a minimum of three individuals in each population, and expected a positive relationship between DMA frequency and *N*_e_.

After determining that there was no evidence of purging, we tested for the expected putative signatures of genetic drift on DMA frequency. To facilitate comparisons among populations that contained different numbers of individuals, for each population we sampled three non-missing genotypes per site (n = 100 sampling replicates). We calculated the mean (across sampling replicates) number of sites with at least one derived allele present, the mean frequency of derived alleles that were present, and the mean fixation rate of derived alleles that were present. We used simple linear regressions to test the expectations that genetic drift would reduce the number of sites with a derived allele present in small populations relative to large populations, and that remaining derived alleles in small populations would have higher mean frequency and be more likely to be fixed. We also estimated the realized load from the resampled datasets as the proportion of genotypes homozygous for the DMA (at evolutionarily conserved sites and moderate- to high-impact sites only) and tested for an expected negative correlation with *N*_e_.

#### Genetic load under different cross types

We examined the effects of selfing on realized genetic load using empirical genetic load estimates from selfed and non-selfed individuals. We first used custom scripts to estimate the realized genetic load for all individuals at putatively deleterious DMAs for both the GERP and *SnpEff* analyses. Using the definitions of which individuals were selfed or non-selfed (described above), we then calculated the relative difference in realized genetic load between each selfed individual and the mean realized genetic load of all non-selfed individuals in the corresponding population. We used linear regressions to test for a relationship between this relative difference in realized load and *N*_e_. Because data points were not independent (i.e., in some cases multiple selfed individuals were compared to the same population mean), we calculated adjusted p-values for the relationship by permuting *N*_e_ 1,000 times and taking the proportion of permuted *t*-statistics that were greater than the empirical *t-*statistic.

We also predicted the effects of different cross types on realized genetic load by comparing simulated crosses within and between populations. We used custom scripts to randomly sample one individual per population for each pairwise combination of populations. For within-population crosses we ensured that first-degree relatives were not sampled and the same individual was not sampled twice, so that we would not simulate any highly inbred or selfed individuals that could confound comparisons. For each sampled pair of individuals, we randomly selected one allele at each site and calculated the realized genetic load of DMAs in the simulated offspring. Sampling of individuals was repeated 100 times for each pair of populations, and we calculated the mean realized load across all replicates of each pair.

To test whether outcrossing between populations reduced realized genetic load relative to crossing within populations, we calculated the relative difference in the mean realized load (across the 100 sampling replicates) of between-population crosses relative to that of within-population crosses for each recipient population. The recipient population was defined as the population used as the comparison in the within-population cross and the donor population as the other population. For example, considering recipient population A and donor population B, the relative difference in realized genetic load was calculated for cross A x B relative to cross A x A. Switching the definition of the recipient and donor populations results in a second comparison of crosses B x A and B x B for the same population pair.

To test our prediction that the reduction in mean realized load upon between-population outcrossing would be greater when the recipient population was small and the donor population was far away, we performed multiple linear regressions using the *lm()* function in *R* v.4.2.3^110^ with the relative realized genetic load as the response variable and the *N*_e_ of the recipient population and the geographic distance between populations as predictor variables. Because the data points were not statistically independent, we calculated adjusted *p*-values by permuting either *N*_e_ or the matrix of geographic distances while holding all other values constant and taking the proportion of permuted *t*-statistics more extreme than the empirical *t*-statistic for the variable of interest. To visualize the three-dimensional relationship between *N*_e_, geographic distance, and relative realized load, we treated the predictor variables as a two-dimensional landscape and performed smoothing and interpolation of the response variable across this landscape using *snapKrig* v.0.0.2^120^, with a grid of 51 x 51 cells for kriging and default parameters to select the maximum likelihood model.

## Acknowledgements

The authors thank M Bartlett, J Baum, N Bercovich, J Braun, H Bregulla, K Bruce, L Chalifour, D Cliffe, L Coleman, J Collens, R Corder, S Crawford, S Dalanson, D Denley, G Fisher, A-M Flores, G Garner, L Gendall, E González, C Harley, S Henderson, F Hernandez, J Lazaro Guevara, S Le Saout, S Lindsay, A McConnell, C Mountain, D Newman, M Norton, C Pfister, W Roberts, T Robinson, S Rogers, C Ryan, A Schubert, J Schuster, E Starr, C Steell, B Sternberg, L Stewart, M Thompson, S Thurber, M van Roy, A Wachmann, A Zielinski, R Zielinski, Cedar Coast Field Station staff and others for project support and sample collection, and the numerous First Nations and Tribes on whose waters and lands this work was conducted. We thank the following First Nations for collaboration through the Marine Plan Partnership for the North Pacific Coast (MaPP): Gitga’at, Gitxaała, Haida, Haisla, Heiltsuk, Kitasoo-Xai’xais, Kitselas, Kitsumkalum, K’ómoks, Mamalilikulla, Metlakatla, Tlowitsis, and Wei Wai Kum. For a Biocultural Notice^72^ regarding samples provided through MaPP, see the Resource Availability section. We also thank the Makah Tribe, the Squaxin Island Tribe, and the Huu-ay-aht, Ka:’yu:’k’t’h’/Che:k:tles7et’h’, Mamalilikulla, ‘Namgis, and Kwikwasut’inuxw Haxwa’mis First Nations for support regarding the collection of kelp from their lands. The giant kelp silhouette was created by Harold N Eyester. Funding was provided by a Genome British Columbia GIRAFF Grant, the Province of British Columbia, the Washington State Legislature 2021-23 proviso for kelp conservation, and donations to The Kelp Rescue Initiative from the Ngan-Page Family Fund. Postdoctoral support was provided by a Mitacs Accelerate Postdoctoral Fellowship, the Pacific Salmon Foundation, and the Forrest Research Foundation. Computational resources were provided by the Canadian Foundation for Innovation, the BC Knowledge Development Fund and the Digital Research Alliance of Canada.

## Author contributions

Conceptualization, LHR, JEP, CJN, GLO; Formal Analysis, JBB, CE; Investigation, JBB, KH, AP; Resources, SS, BLW, CJN; Writing – Original Draft, JBB; Writing – Review & Editing, all authors; Supervision, MND, LHR, GLO; Project Administration, CJN; Funding Acquisition, MND, LHR, JEP, CJN, GLO.

## Resource availability

- Raw DNA sequences generated in this study have been deposited at NCBI and will be released in the future once this preprint is accepted for publication. Some of the raw DNA sequences are derived from samples (identified in Table S1) subject to a Biocultural Notice^72^: “The BC (Biocultural) Notice is a visible notification that there are accompanying cultural rights and responsibilities that need further attention for any future sharing and use of this material or data. The BC Notice recognizes the rights of Indigenous peoples to permission the use of information, collections, data and digital sequence information (DSI) generated from the biodiversity or genetic resources associated with traditional lands, waters, and territories. The BC Notice may indicate that BC Labels are in development and their implementation is being negotiated. For more information about the BC Notice see <u>localcontexts.org/notice/bc-notice/</u>.”
- All original code will be deposited at Zenodo in the future once this preprint is accepted for publication.
- Any additional information required to reanalyze the data reported in this paper will be available upon request in the future once this preprint is accepted for publication.

## Supplemental info

Supplemental information (Figures S1-S12 and Tables S1-S4) is available with the online version of this preprint.

