## Supplemental information for "Population genomics reveals strong impacts of genetic drift without purging and guides conservation of bull and giant kelp"

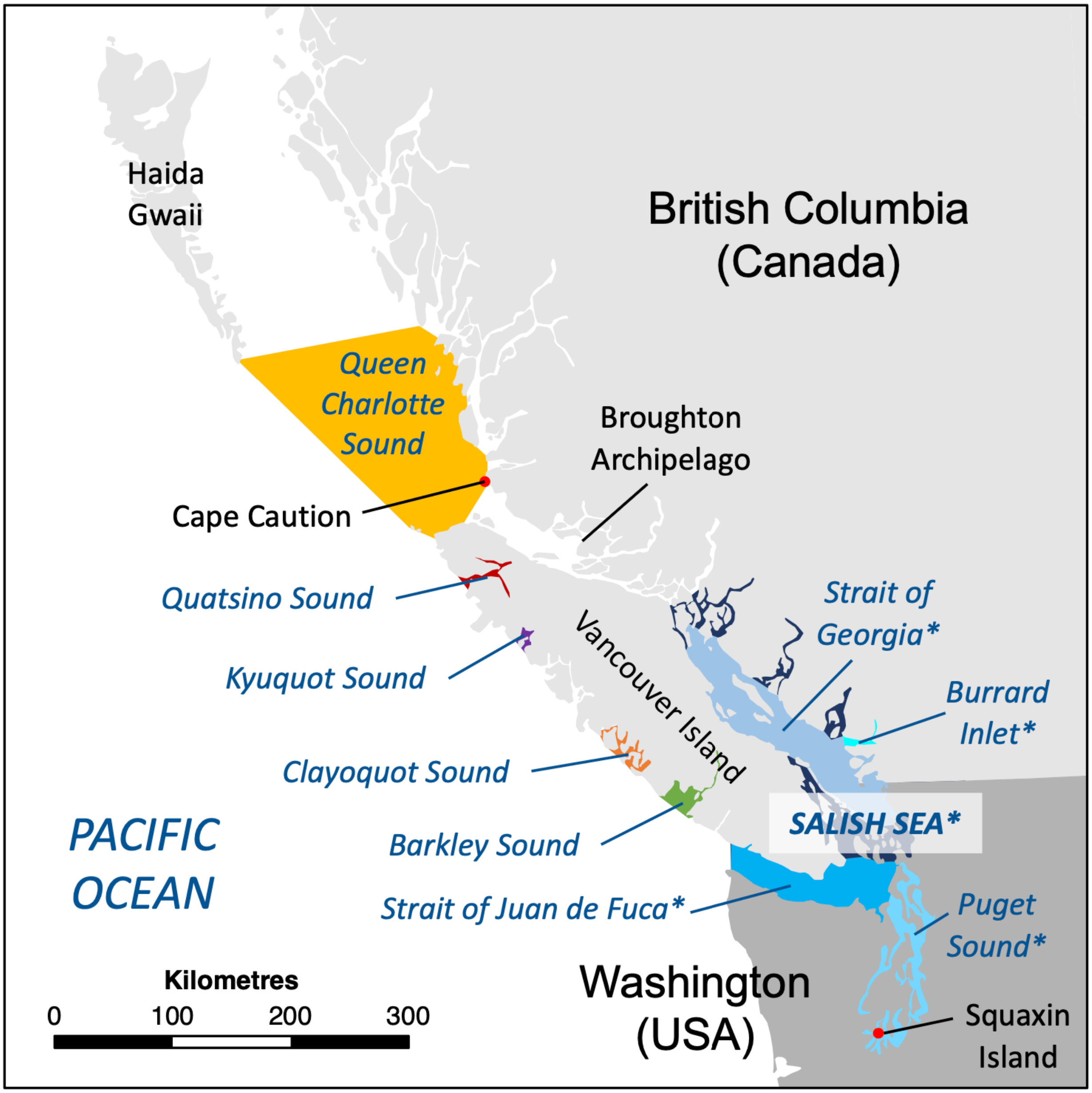


**Figure S1. Map of geographic place names, Related to Figure 1**
Map of geographic place names mentioned in this paper. Terrestrial features are denoted in black text and oceanic features in italicized blue text. The Salish Sea includes all areas in various shades of blue, including separately labelled subregions of the Salish Sea indicated with an asterisk (*). Boundaries of all oceanic features are approximate.

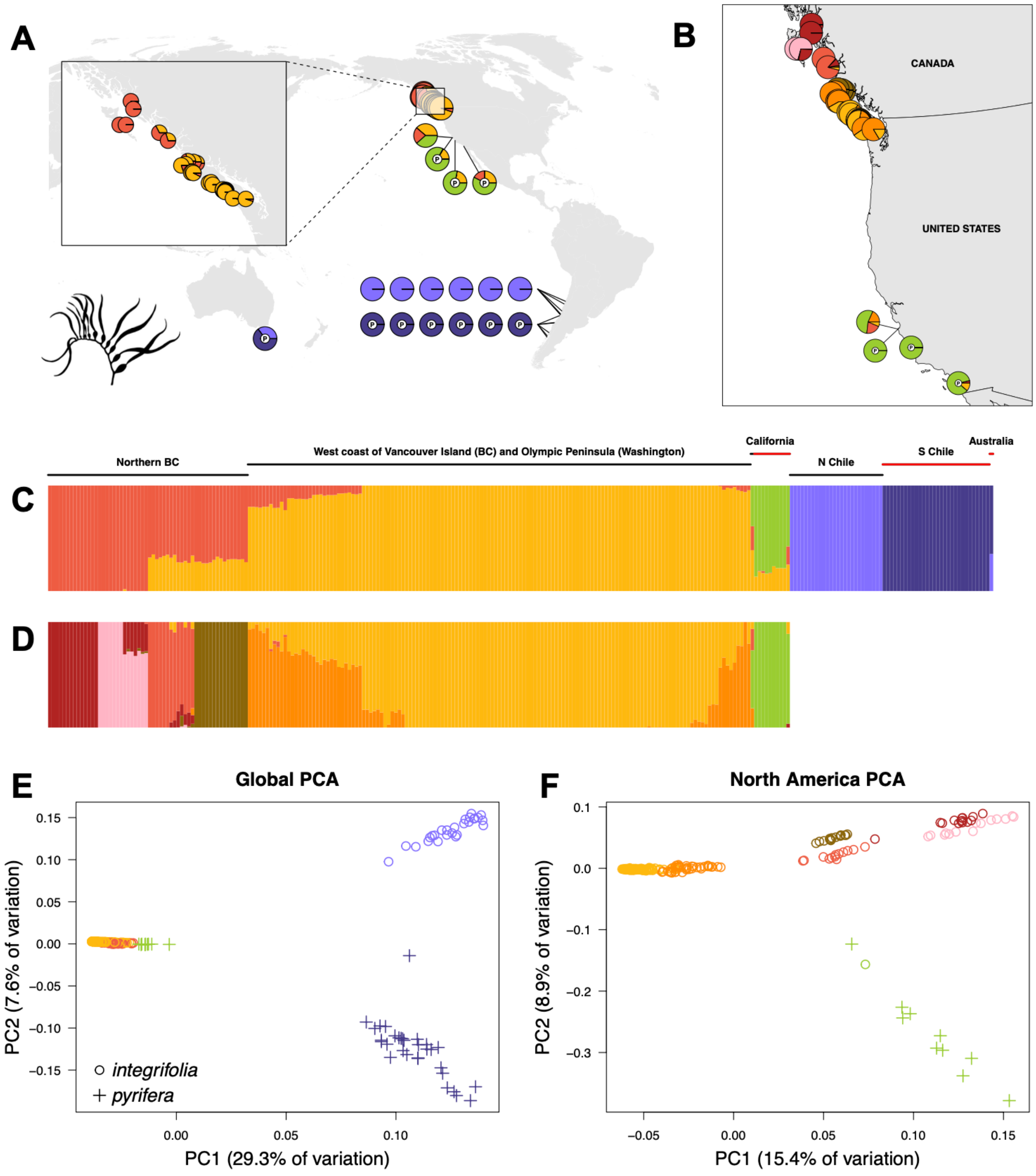


**Figure S2. Global genetic structure of giant kelp, Related to Figure 1**
Genetic structure estimated from (A, C, E) global samples and (B, D, F) North American samples only. (A-B) Pie charts represent the proportion of ancestry in each population belonging to different genetic clusters, with clusters represented as different colours. (C-D) The proportion of ancestry derived from each genetic cluster in each individual, with individuals represented as vertical lines. (E-F) Principal component analysis showing the clustering of individuals along the first two PC axes. Each of the clusters from (A-D) is represented by a different symbol and its corresponding colour. Samples of the *pyrifera* ecomorph have a small “P” in the middle of each pie chart in (A-B), are outlined in red above the individuals in (C-D), and are indicated by a (+) symbol in (E-F); otherwise samples are of the *integrifolia* ecomorph.

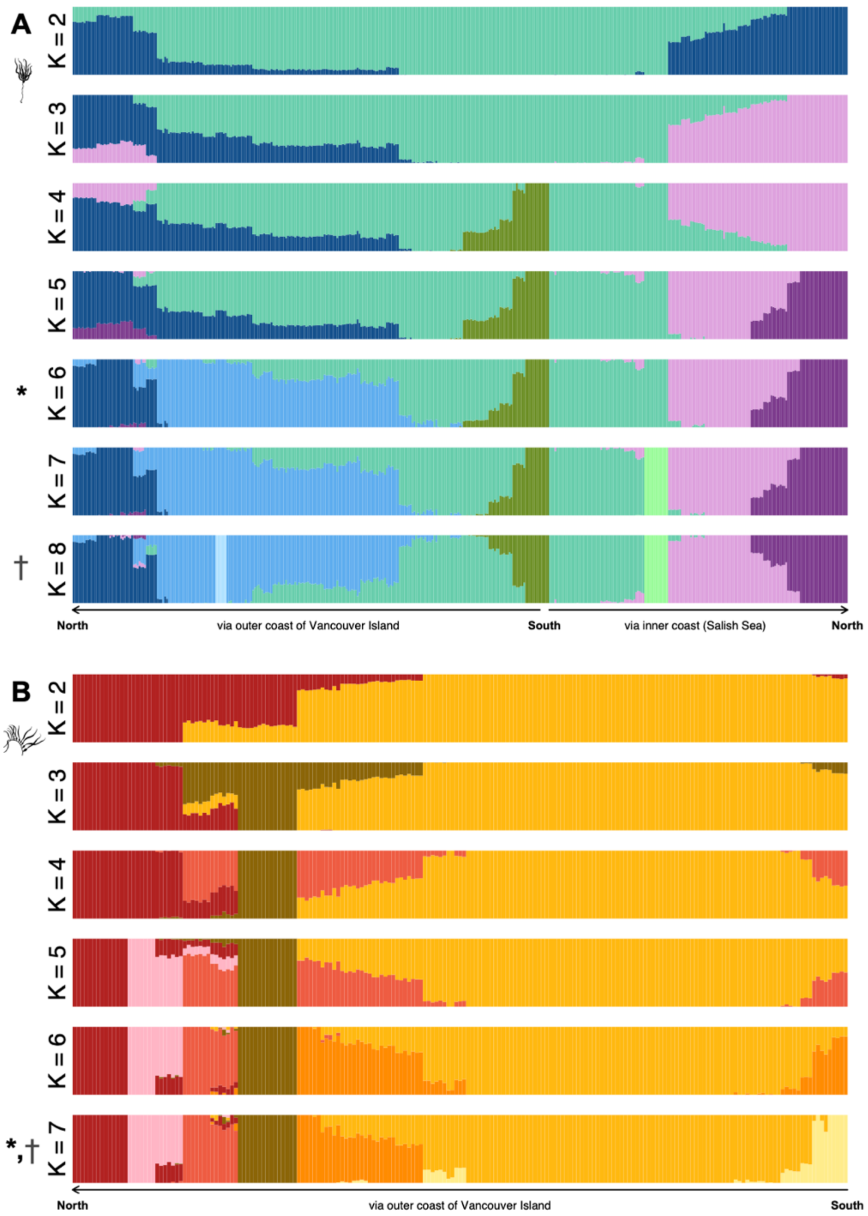


**Figure S3. Hierarchical genetic clustering, Related to Figure 1**
Genetic structure across British Columbia and Washington with different number of clusters (K) in (A) bull kelp and (B) giant kelp. Colours represent the proportion of ancestry derived from each genetic cluster in each individual, with individuals represented as vertical lines. Asterisk (*), main model selected for presentation in Figure 1 and throughout this manuscript; dagger (†), statistically optimal number of clusters (see STAR Methods). In bull kelp, K = 6 was selected as the main model in the manuscript despite K = 8 being the statistically optimal model because the two additional clusters identified in K = 8 represent very small geographic areas where individual or closely-related populations are genetically depauperate relative to neighbouring populations (Figure 2) and may have experienced recent bottlenecks, rather than broader-scale clusters that reflect regional genetic patterns. These clusters are located in Burrard Inlet near Vancouver (brightest green, K = 7 and K = 8) and an isolated innermost area of Clayoquot Sound on the west coast of Vancouver Island (brightest blue, K = 8).

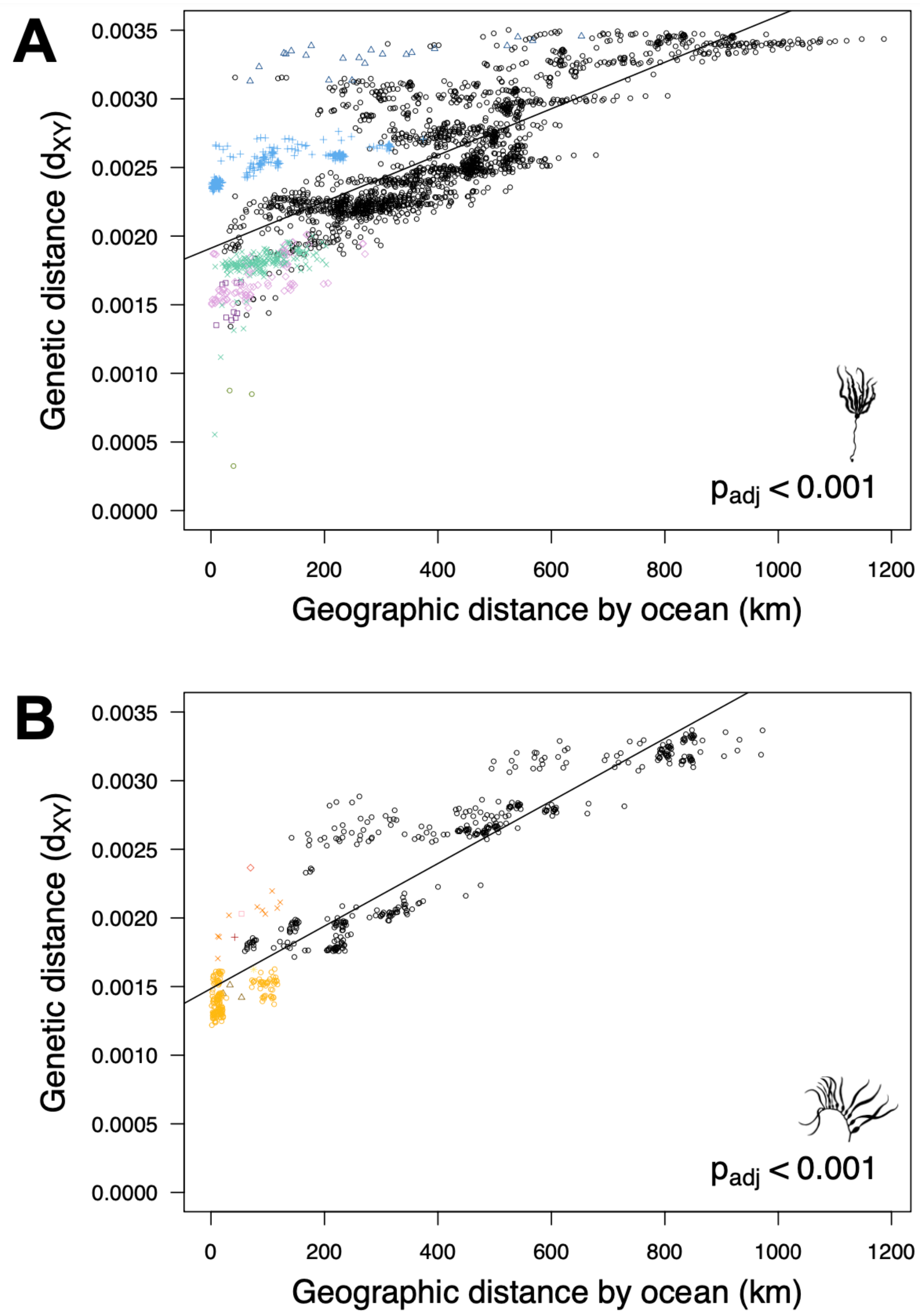


**Figure S4. Isolation by distance pattern, Related to Figure 1**
Genetic distance ($d_{XY}$) versus geographic distance by the shortest ocean route for (A) bull kelp and (B) giant kelp. Each point represents a pair of populations. Comparisons within the same genetic cluster are represented with the same colours and symbols for each cluster as in Figure 1, while comparisons between different genetic clusters are represented as black open circles. Adjusted p-values are from 1,000 permutations of the geographic distance matrix.

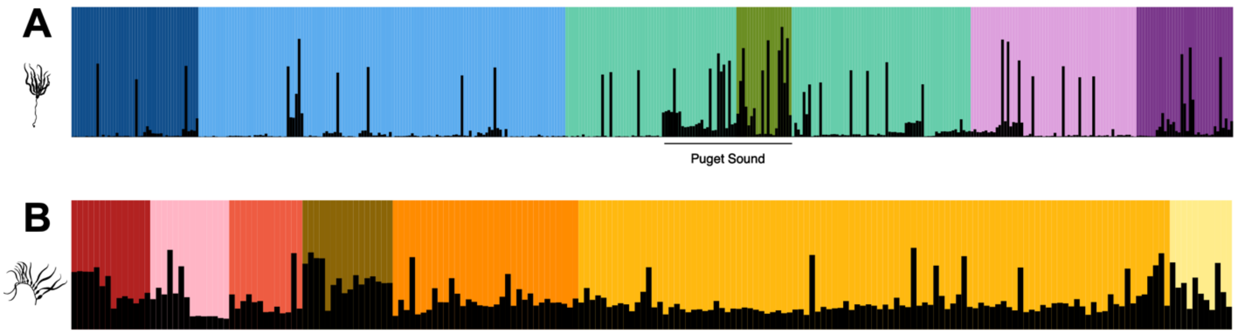


**Figure S5. Runs of homozygosity, Related to Figure 2**
Runs of homozygosity (ROHs) ≥100 kbp in length in each individual of (A) bull kelp and (B) giant kelp. Each vertical bar represents an individual, with the colour of the bar representing the genetic cluster from Figure 1 for which the individual has the largest proportion of ancestry. The black portion of each bar represents the proportion of each individual's genome in ROHs, with a higher proportion representing a higher inbreeding coefficient ($F_{ROH\_100kbp}$). The location of Puget Sound, Washington is highlighted in (A) to indicate populations with elevated inbreeding coefficients in bull kelp that span two different genetic clusters but represent a contiguous geographic region.

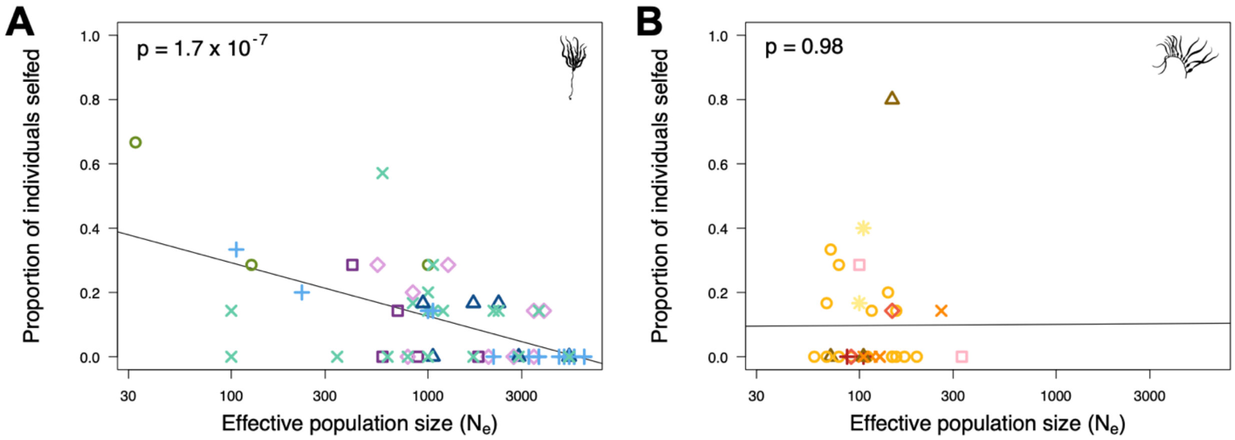


**Figure S6. Relationship between selfing rate and population size, Related to Figure 2**
Selfing rate (empirical proportion of individuals assumed to be selfed) versus effective population size ($N_e$) for (A) bull kelp and (B) giant kelp. Each point represents a population. Symbols and colours correspond to genetic clusters from Figure 1.

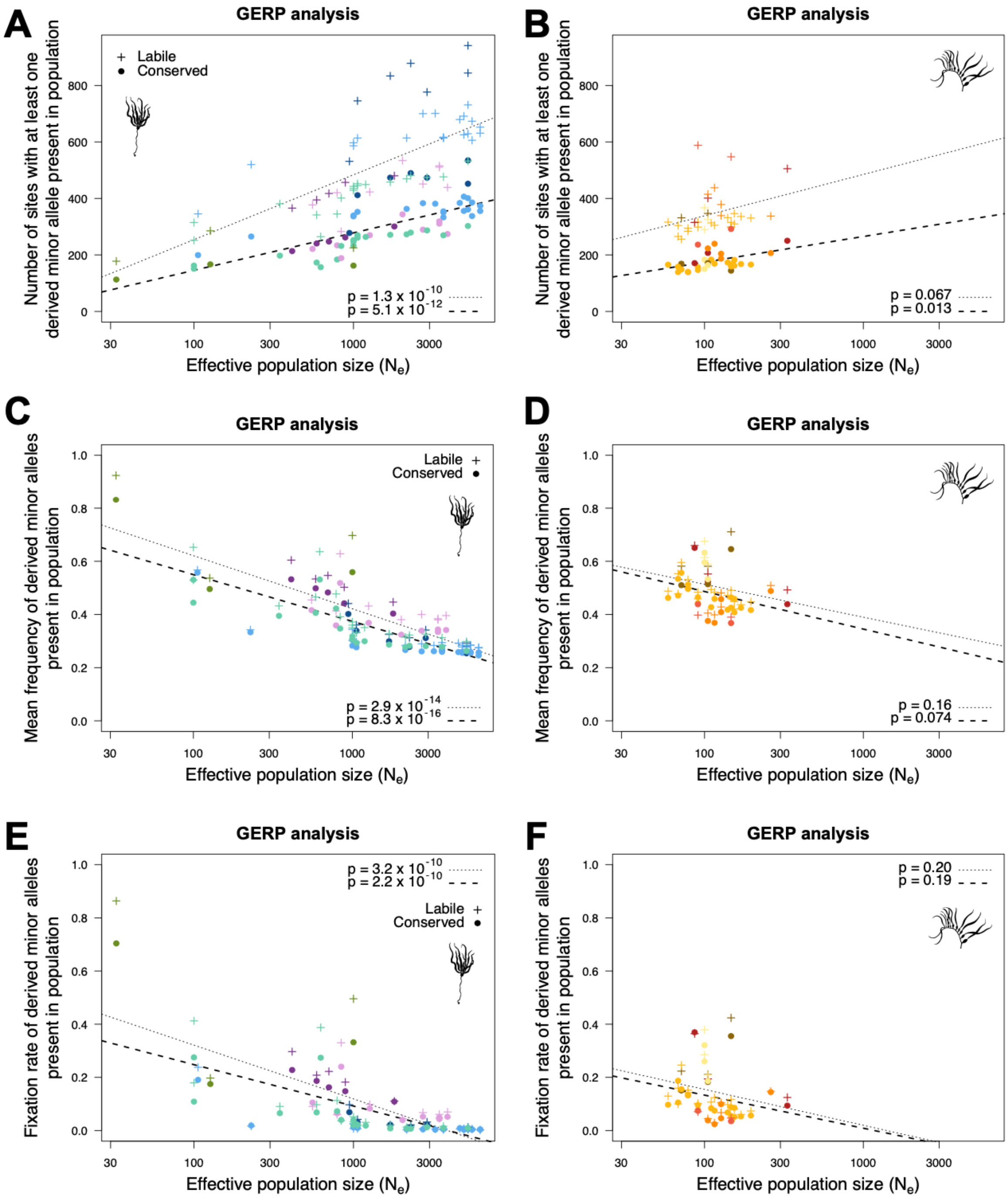


**Figure S7. Effects of genetic drift from GERP analysis, Related to Figure 4**
Tests for shifts in allele frequency of evolutionarily labile and conserved sites in small populations of (A, C, E) bull kelp and (B, D, F) giant kelp. (A-B) Number of sites with at least one derived minor allele (DMA) present in at least one individual in the population versus effective population size ($N_e$). (C-D) Mean frequency of DMAs that are present in the population versus $N_e$. (E-F) Proportion of DMAs present in the population that are fixed versus $N_e$. Each point represents a population. Symbols and colours correspond to genetic clusters from Figure 1. In all panels, populations were resampled to n = 3 individuals to facilitate direct comparison between populations of different sample size.

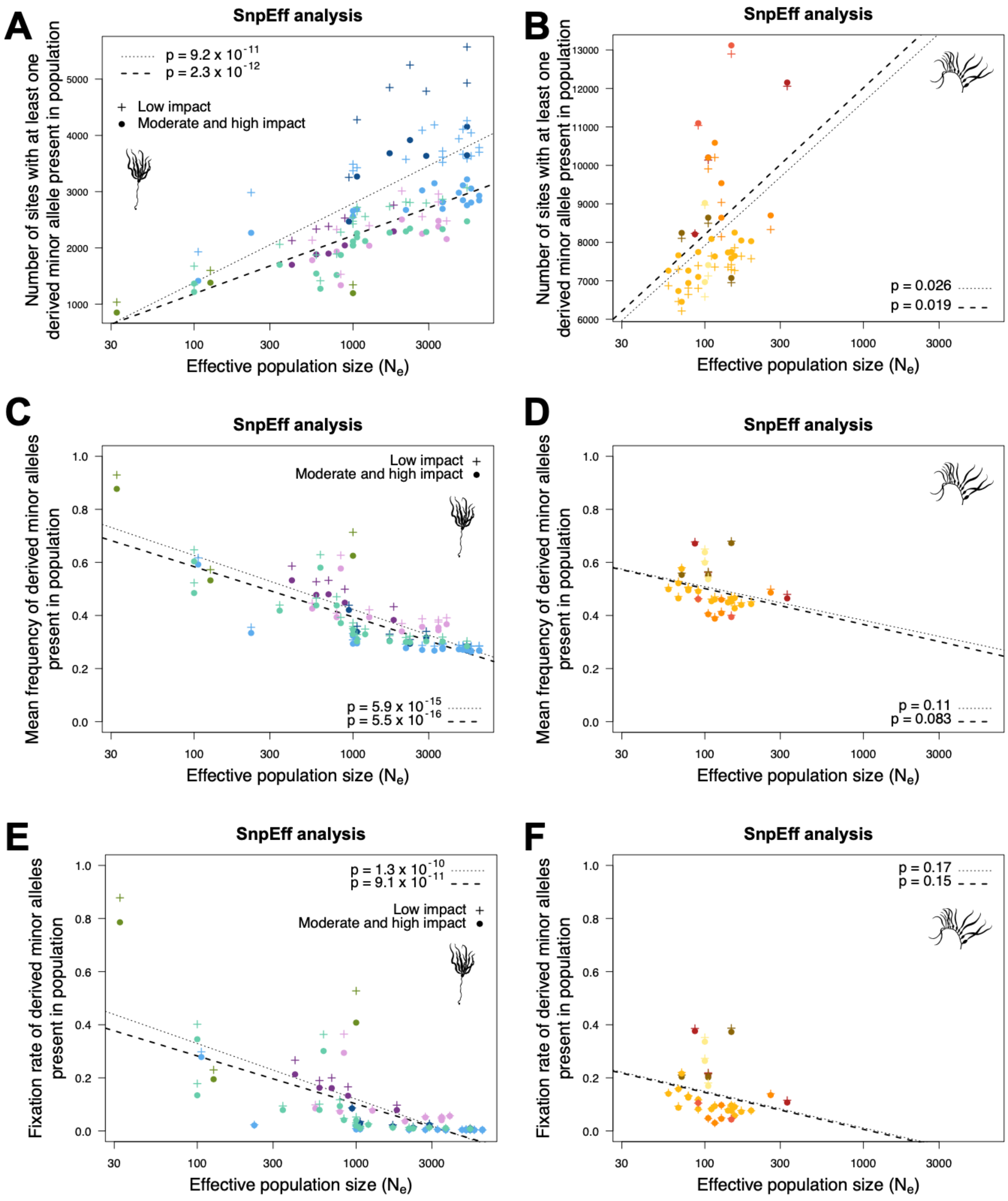


**Figure S8. Effects of genetic drift from *SnpEff* analysis, Related to Figure 4**

Tests for shifts in allele frequency of low impact and moderate- to high-impact sites in small populations of (A, C, E) bull kelp and (B, D, F) giant kelp. (A-B) Number of sites with at least one derived minor allele (DMA) present in at least one individual in the population versus effective population size ($N_e$). (C-D) Mean frequency of DMAs that are present in the population versus $N_e$. (E-F) Proportion of DMAs present in the population that are fixed versus $N_e$. Each point represents a population. Symbols and colours correspond to genetic clusters from Figure 1. In all panels, populations were resampled to n = 3 individuals to facilitate direct comparison between populations of different sample size.

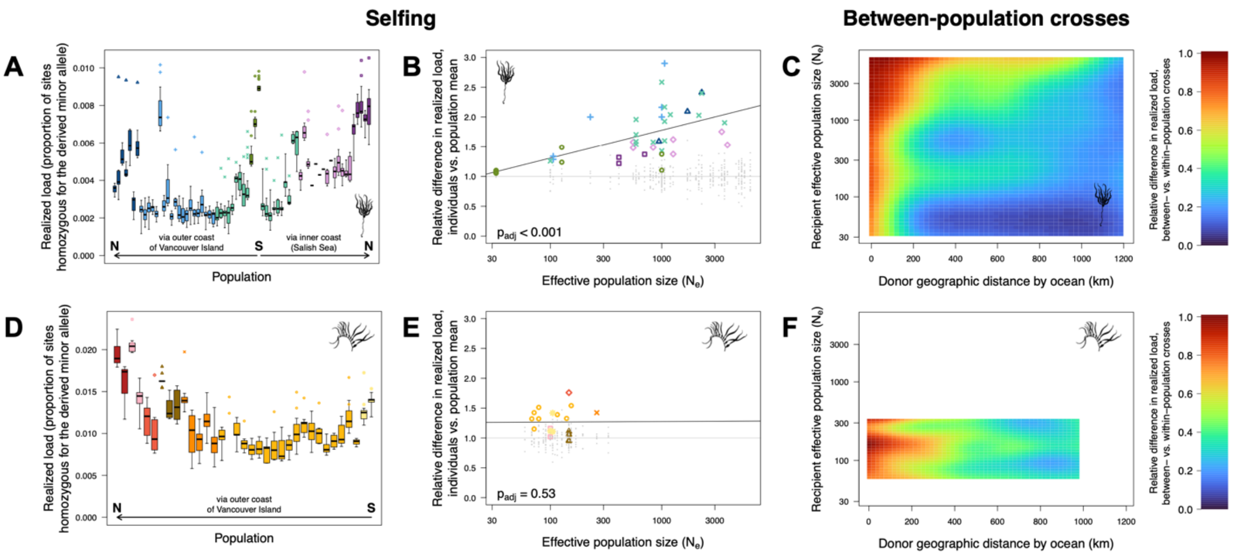


**Figure S9. Predicted genetic load under different cross types, GERP analysis, Related to Figure 4**
Predicted changes in realized genetic load under different crosses for (A-C) bull kelp and (D-F) giant kelp result in (A-B, D-E) a predicted penalty for selfing and (C, F) predicted heterosis in between-population crosses. Realized genetic load is measured as the proportion of sites homozygous for the derived allele at evolutionarily conserved sites in the GERP analysis. (A, D) Realized load is higher in selfed individuals (point symbols, one point per individual) than in non-selfed individuals (distribution in boxplots, with boxplot whiskers extending to the most extreme values). Symbols and colours correspond to genetic clusters from Figure 1. (B, E) Relative difference in realized load of selfed individuals (coloured symbols, one point per individual) compared to the mean realized load of non-selfed individuals versus effective population size ($N_e$). Non-selfed individuals are shown in small grey points for comparison. Adjusted p-values are calculated from 1,000 randomizations of $N_e$. (C, F) Realized genetic load is predicted to be equal or lower (≤1.0) in between-population crosses (recipient x donor population) relative to within-population crosses (recipient x recipient). The predicted relative load (colour scale) is plotted as a function of recipient $N_e$ and geographic distance by ocean to the donor population. The coloured two-dimensional surface is inferred by kriging of raw data points (Figure S11).

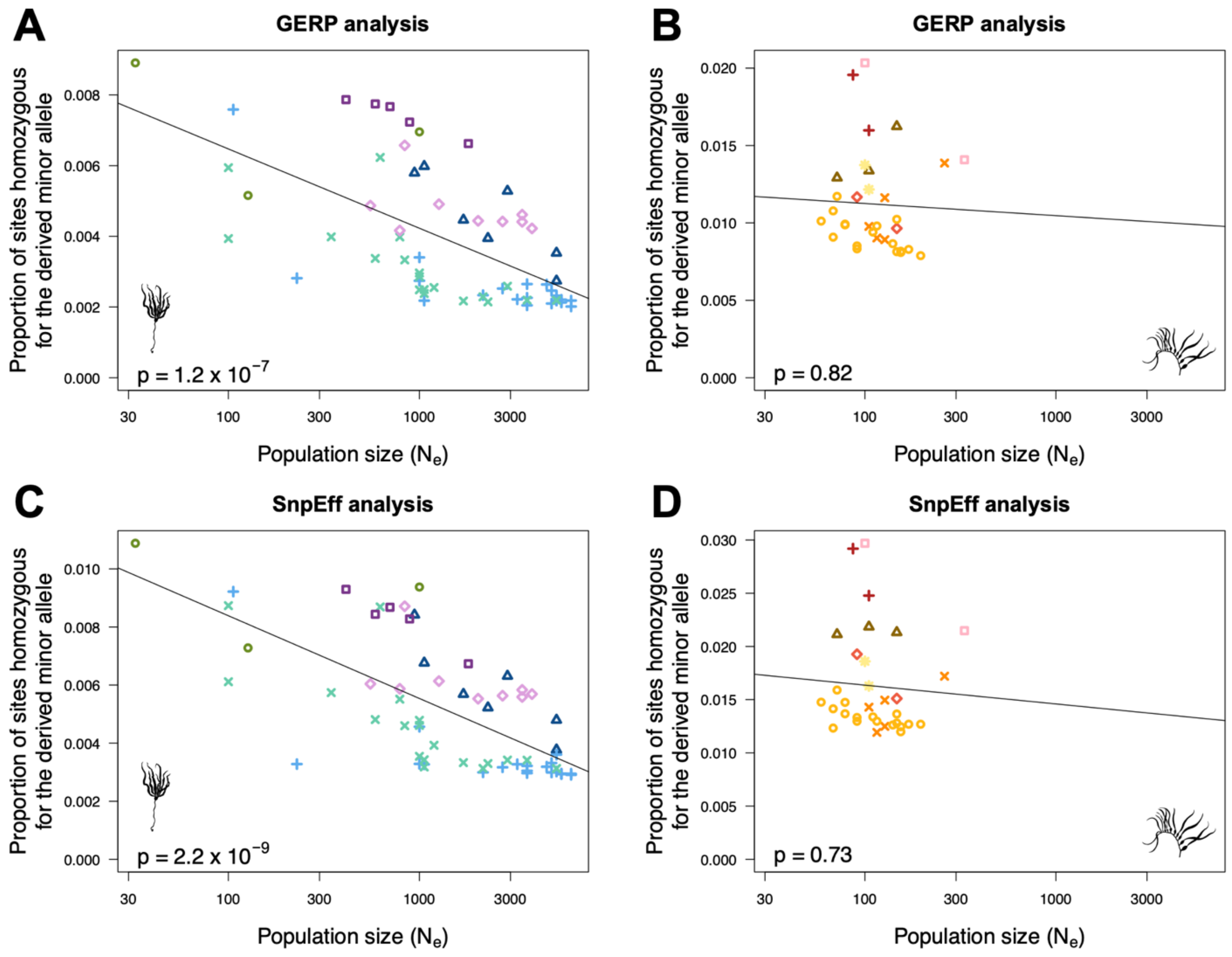


**Figure S10. Relationship between realized load and population size, Related to Figure 4**
Relationship between realized genetic load, measured as the proportion of sites homozygous for the derived minor allele in deleterious allele categories, and effective population size ($N_e$). Realized load is calculated from non-selfed individuals only. Relationships are shown for (A, C) bull kelp and (B, D) giant kelp. (A-B) In GERP analyses, realized load is calculated from evolutionarily conserved sites. (C-D) In *SnpEff* analyses, realized load is calculated from moderate- and high-impact sites. Each point represents a population. Symbols and colours correspond to genetic clusters from Figure 1.

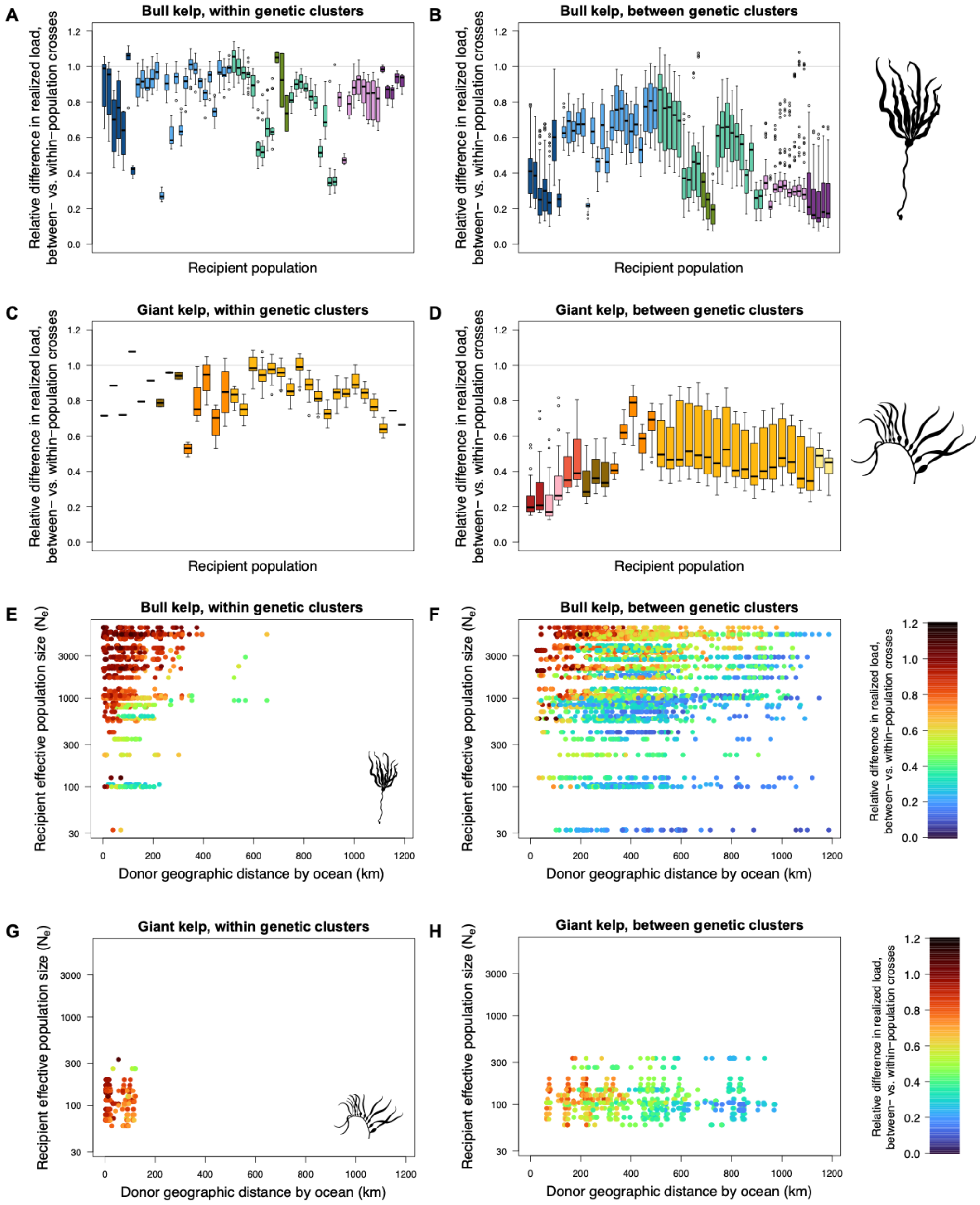


**Figure S11. Realized load in between-population crosses (GERP), Related to Figure 4**
Relative difference in realized genetic load in simulated between-population crosses (recipient x donor population) relative to simulated within-population crosses (recipient x recipient) for (A-B, E-F) bull kelp and (C-D, G-H) giant kelp. Realized genetic load is measured as the proportion of sites homozygous for the derived allele at evolutionarily conserved sites in the GERP analysis. (A-D) Boxplots of the distribution of relative realized genetic load across all possible donor populations for each recipient population, when the donor population is sourced (A, C) within or (B, D) between genetic clusters. Colours correspond to genetic clusters from Figure 1. (E-H) Effects of recipient effective population size ($N_e$) and geographic distance by ocean to the donor population (km) on the relative realized genetic load, when the donor population is sourced (E, G) within or (F, H) between genetic clusters. Each dot represents a pair of recipient and donor populations, with the colours indicating the realized genetic load relative to the corresponding within-population (recipient x recipient) cross.

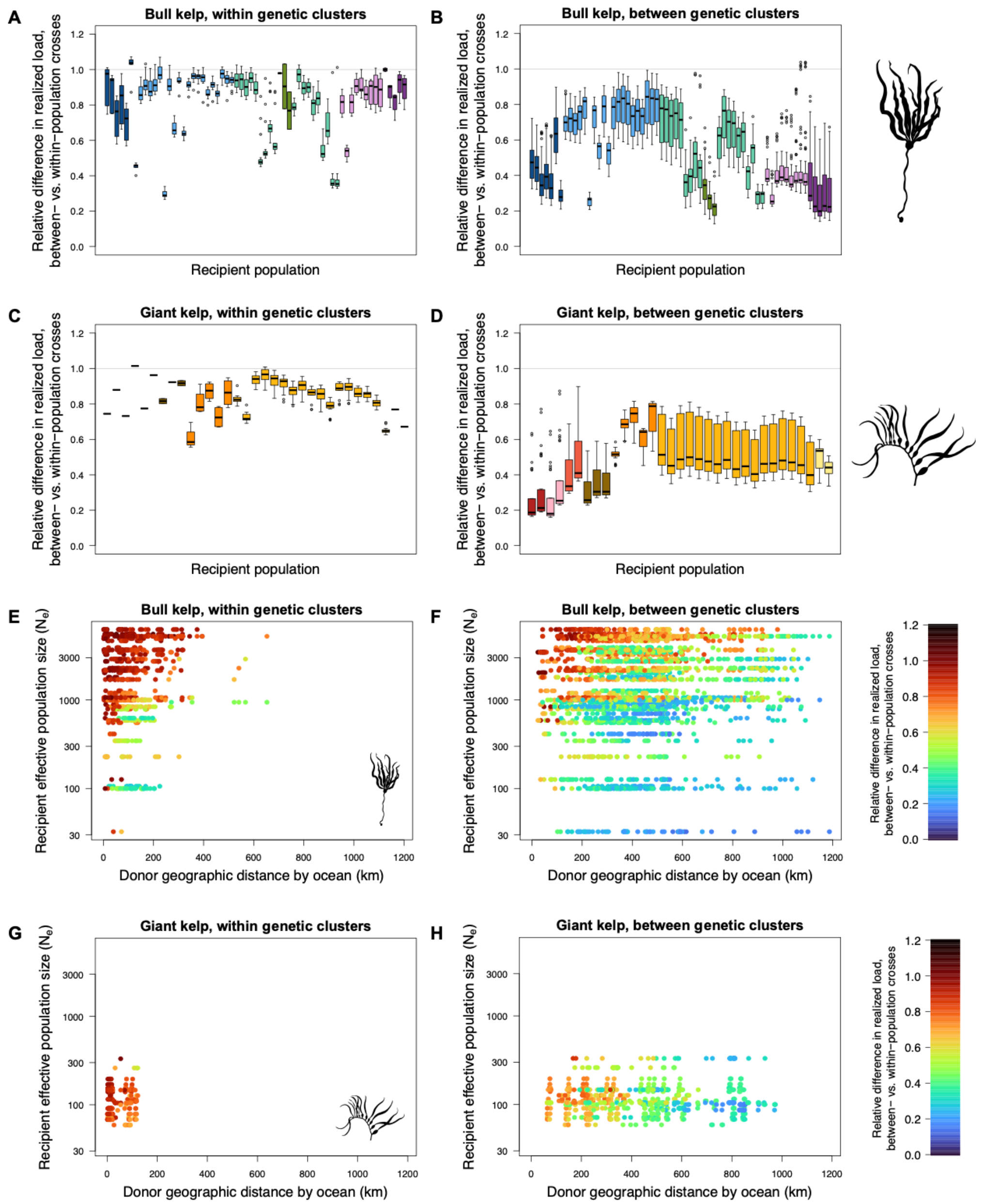


**Figure S12. Realized load in between-population crosses (*SnpEff*), Related to Figure 4**
Relative difference in realized genetic load in simulated between-population crosses (recipient x donor population) relative to simulated within-population crosses (recipient x recipient) for (A-B, E-F) bull kelp and (C-D, G-H) giant kelp. Realized genetic load is measured as the proportion of sites homozygous for the derived allele at moderate- and high-impact sites in the *SnpEff* analysis. (A-D) Boxplots of the distribution of relative realized genetic load across all possible donor populations for each recipient population, when the donor population is sourced (A, C) within or (B, D) between genetic clusters. Colours correspond to genetic clusters from Figure 1. (E-H) Effects of recipient effective population size ($N_e$) and geographic distance by ocean to the donor population (km) on the relative realized genetic load, when the donor population is sourced (E, G) within or (F, H) between genetic clusters. Each dot represents a pair of recipient and donor populations, with the colours indicating the realized genetic load relative to the corresponding within-population (recipient x recipient) cross.

**Table S1. Sampling information, related to Figure 1.**
Sampling sites and sample sizes for (A) bull kelp and (B) giant kelp. Abbreviations for political jurisdictions (Jur.) are as follows: AU, Australia; BC, British Columbia; CA, California; CL, Chile; WA, Washington. Sample sizes refer to the number of individuals retained at different stages of the bioinformatics pipeline: $n_{seq}$, total number of individuals sequenced; $n_{dep}$, number of individuals for which sufficient sequencing depth was obtained (see STAR Methods); $n_{uni}$, number of unique individuals retained after removing genetically identical samples, $n_{unr}$, number of unrelated individuals retained after additionally removing putative first-degree relatives. Samples corresponding to sites and coordinates marked with an asterisk (*) or dagger (†) are subject to a Biocultural Notice[72] (see the Resource Availability section), with First Nations providing specific samples indicated in the site name; NC Nations: Gitga'at, Gitxaała, Haisla, Kitselas, Kitsumkalum, and Metlakatla First Nations. (*) Latitude and longitude rounded to the nearest 0.5º. (†) Latitude and longitude rounded to the nearest 0.1º. (‡) Precise coordinates are unknown; approximate coordinates are provided and were obtained by georeferencing the site name. (§) Samples are unlikely to represent populations native to the collecting site and were removed from most genetic analyses.

*(Table displayed on following page)*

| Jur. | Site code | Site name | Latitude | Longitude | $n_{seq}$ | $n_{dep}$ | $n_{uni}$ | $n_{unr}$ | Citation |
|---|---|---|---:|---:|---|---|---|---|---|
| BC | NL-HG-03 | [*] Haida 03 | 54.0[*] | -132.0[*] | 7 | 7 | 7 | 7 | this study |
| BC | NL-HG-01 | [*] Haida 01 | 53.0[*] | -131.5[*] | 7 | 7 | 6 | 6 | this study |
| BC | NL-NC-02 | [*] NC Nations 02 | 54.5[*] | -130.5[*] | 7 | 7 | 7 | 7 | this study |
| BC | NL-NC-01 | [*,‡] NC Nations 01 | 54.0[*,‡] | -130.5[*,‡] | 7 | 7 | 6 | 6 | this study |
| BC | NL-CC-03 | [*] Kitasoo-Xai'xais 03 | 52.5[*] | -128.5[*] | 7 | 7 | 7 | 7 | this study |
| BC | NL-CC-02 | [*] Heiltsuk 02 | 52.0[*] | -128.5[*] | 7 | 7 | 7 | 7 | this study |
| BC | NL-QS-07 | Kenny Point | 50.56003 | -127.53754 | 7 | 6 | 6 | 6 | this study |
| BC | NL-QS-01 | Kains Island | 50.44589 | -128.03422 | 7 | 7 | 6 | 6 | this study |
| BC | NL-KQ-01 | SE Spring Island | 50.00193 | -127.43080 | 7 | 7 | 5 | 5 | this study |
| BC | NL-KQ-03 | Duchess Cove | 50.01231 | -127.32945 | 7 | 7 | 7 | 7 | this study |
| BC | NL-KQ-05 | Rugged Point | 49.97070 | -127.25286 | 7 | 7 | 7 | 7 | this study |
| BC | NL-CS-03 | Maquinna Park | 49.39444 | -126.34806 | 7 | 7 | 7 | 7 | this study |
| BC | NL-CS-02 | Obstruction Island | 49.41472 | -126.06861 | 7 | 6 | 6 | 6 | this study |
| BC | NL-CS-06 | [‡] Catface | 49.24061[‡] | -126.01013[‡] | 7 | 7 | 7 | 7 | this study |
| BC | NL-CS-07 | [‡] Matlset | 49.23719[‡] | -125.79528[‡] | 7 | 7 | 7 | 7 | this study |
| BC | NL-BS-01 | Clarke Island | 48.88944 | -125.37241 | 7 | 7 | 6 | 6 | this study |
| BC | NL-BS-05 | Mullins Island | 48.90728 | -125.30055 | 7 | 5 | 5 | 5 | this study |
| BC | NL-BS-11 | Folger | 48.83083 | -125.24803 | 7 | 7 | 7 | 7 | this study |
| BC | NL-BS-12 | Bordelais | 48.81814 | -125.22970 | 7 | 7 | 7 | 7 | this study |
| BC | NL-BS-13 | Skull Bay | 48.82637 | -125.22238 | 7 | 7 | 7 | 7 | this study |
| BC | NL-BS-14 | BBNEK | 48.82727 | -125.21169 | 7 | 7 | 7 | 7 | this study |
| BC | NL-BS-16 | Helby | 48.85527 | -125.16855 | 7 | 7 | 7 | 6 | this study |
| BC | NL-BS-24 | Aguilar | 48.83945 | -125.14094 | 7 | 7 | 7 | 7 | this study |
| BC | NL-BS-19 | Ross Three-Tree Island | 48.87463 | -125.16407 | 7 | 7 | 7 | 7 | this study |
| BC | NL-BS-20 | Tzartus Backside | 48.90692 | -125.11360 | 7 | 7 | 7 | 7 | this study |
| BC | NL-BS-22 | Island by Mudd Cove | 48.79347 | -125.21465 | 7 | 7 | 7 | 7 | this study |
| WA | NL-WC-01 | Tatoosh Island | 48.39361 | -124.73515 | 7 | 7 | 7 | 7 | this study |
| WA | NL-WC-02 | Clallam Bay | 48.25633 | -124.27249 | 7 | 7 | 7 | 7 | this study |
| BC | NL-SK-01 | Ella Beach | 48.35709 | -123.74846 | 7 | 7 | 7 | 7 | this study |
| WA | NL-WC-03 | Freshwater Bay | 48.14944 | -123.63806 | 7 | 7 | 7 | 7 | this study |
| BC | NL-VI-01 | Ogden Point | 48.41370 | -123.38697 | 7 | 7 | 7 | 7 | this study |
| WA | NL-WC-04 | Fort Worden State Park | 48.14384 | -122.76414 | 7 | 7 | 7 | 7 | this study |
| WA | NL-PS-01 | Polnell Point | 48.27103 | -122.55694 | 7 | 7 | 7 | 7 | this study |
| WA | NL-PS-02 | Camano Island State Park | 48.12897 | -122.50619 | 7 | 7 | 7 | 7 | this study |
| WA | NL-PS-03 | Hansville | 47.92013 | -122.55522 | 7 | 6 | 6 | 6 | this study |
| WA | NL-PS-04 | Edmonds | 47.82226 | -122.37611 | 7 | 7 | 7 | 7 | this study |
| WA | NL-PS-05 | Lincoln Park | 47.53339 | -122.39880 | 7 | 7 | 7 | 7 | this study |
| WA | NL-PS-06 | Salmon Beach | 47.29780 | -122.53523 | 7 | 7 | 7 | 7 | this study |
| WA | NL-PS-07 | Squaxin Island | 47.17740 | -122.91171 | 7 | 7 | 6 | 6 | this study |
| WA | NL-SJ-02 | Deadman Bay | 48.51285 | -123.14618 | 7 | 7 | 6 | 6 | this study |
| WA | NL-SJ-04 | Harbor Rock | 48.469 | -122.969 | 7 | 7 | 7 | 7 | this study |
| WA | NL-SJ-03 | Turn Rock | 48.535 | -122.964 | 7 | 7 | 7 | 7 | this study |
| WA | NL-BE-02 | Cababa Park | 48.49098 | -122.69048 | 7 | 7 | 7 | 7 | this study |
| WA | NL-SJ-01 | Gordon Island | 48.73 | -123.021 | 7 | 7 | 7 | 7 | this study |
| WA | NL-BE-01 | Cherry Point | 48.85295 | -122.72616 | 7 | 7 | 7 | 7 | this study |
| BC | NL-CW-01 | Genoa Bay | 48.75766 | -123.59377 | 7 | 7 | 6 | 6 | this study |
| BC | NL-NN-01 | False Narrows | 49.13665 | -123.78818 | 7 | 6 | 5 | 5 | this study |
| BC | NL-VR-01 | Brockton Point | 49.30086 | -123.12187 | 7 | 7 | 7 | 7 | this study |
| BC | NL-VR-02 | New Brighton Park | 49.29108 | -123.03824 | 7 | 6 | 6 | 6 | this study |
| BC | NL-HY-03 | [§] Deep Bay (drift kelp) | 49.45775[§] | -124.73345[§] | 1 | 1 | 1 | 1 | this study |
| BC | NL-HY-01 | [§] Maude Reef (attached) | 49.49897[§] | -124.68186[§] | 1 | 1 | 1 | 1 | this study |
| BC | NL-HY-02 | [§] Maude Reef (drift kelp) | 49.49897[§] | -124.68186[§] | 1 | 1 | 1 | 1 | this study |
| BC | NL-CR-02 | Oyster River Estuary | 49.87070 | -125.11015 | 7 | 7 | 7 | 7 | this study |
| BC | NL-SC-01 | Skookumchuck Narrows | 49.73938 | -123.89357 | 7 | 7 | 5 | 5 | this study |
| BC | NL-CR-01 | Brown's Bay | 50.16282 | -125.37443 | 7 | 7 | 7 | 7 | this study |
| BC | NL-CR-03 | Whiterock Passage | 50.24775 | -125.10363 | 1 | 1 | 1 | 1 | this study |
| BC | NL-CR-04 | Hole in the Wall | 50.29977 | -125.20895 | 1 | 1 | 1 | 1 | this study |
| BC | NL-CR-05 | Diamond Bay | 50.30217 | -125.22220 | 1 | 1 | 1 | 1 | this study |
| BC | NL-CR-06 | Owen Bay | 50.31107 | -125.22860 | 1 | 1 | 1 | 1 | this study |
| BC | NL-NV-03 | [†] Wei Wai Kum 03 | 50.3[†] | -125.3[†] | 7 | 7 | 7 | 7 | this study |
| BC | NL-CR-07 | Dent Rapids | 50.41153 | -125.22052 | 1 | 1 | 1 | 1 | this study |
| BC | NL-SW-01 | Kelsey Bay | 50.39717 | -125.96057 | 7 | 7 | 7 | 7 | this study |
| BC | NL-NV-04 | [†] K'ómoks 04 | 50.5[†] | -126.0[†] | 7 | 7 | 7 | 7 | this study |
| BC | NL-NV-02 | [†] Mamalilikulla 02 | 50.6[†] | -126.8[†] | 7 | 7 | 7 | 7 | this study |
| BC | NL-BA-01 | Blowhole | 50.58847 | -126.75933 | 7 | 7 | 6 | 6 | this study |
| BC | NL-BA-03 | Bold Head | 50.62152 | -126.74211 | 7 | 7 | 7 | 7 | this study |
| BC | NL-NV-01 | [†] Tlowitsis 01 | 50.6[†] | -126.2[†] | 7 | 7 | 7 | 7 | this study |
| BC | NL-BA-04 | Gilford Point | 50.65511 | -126.43866 | 7 | 7 | 7 | 6 | this study |

| | | | | | | | | | |
|---|---|---|---|---|---|---|---|---|---|
| BC | NL-BA-06 | Shelterless Point | 50.67423 | -126.09747 | 7 | 7 | 7 | 6 | this study |
| BC | NL-BA-05 | Smith Rock | 50.80808 | -126.42172 | 7 | 7 | 7 | 7 | this study |
| BC | NL-BA-08 | Glacier Falls | 50.84881 | -126.32331 | 7 | 7 | 7 | 7 | this study |
| | | | | **Total** | **449** | **442** | **429** | **426** | |

| **Jur.** | **Site Code** | **Site name** | **Latitude** | **Longitude** | $n_{seq}$ | $n_{dep}$ | $n_{uni}$ | $n_{unr}$ | **Citation** |
|---|---|---|---|---|---|---|---|---|---|
| BC | MP-NC-02 | * NC Nations 02 | 54.5* | -130.5* | 7 | 7 | 7 | 7 | this study |
| BC | MP-NC-01 | * NC Nations 01 | 54.0* | -130.5* | 7 | 7 | 7 | 7 | this study |
| BC | MP-HG-02 | * Haida 02 | 53.0* | -132.0* | 7 | 7 | 7 | 7 | this study |
| BC | MP-HG-01 | * Haida 01 | 53.0* | -131.5* | 7 | 7 | 7 | 7 | this study |
| BC | MP-CC-05 | * Kitasoo-Xai'xais 05 | 52.5* | -129.0* | 7 | 6 | 6 | 6 | this study |
| BC | MP-CC-01 | * Heiltsuk 01 | 52.0* | -128.5* | 7 | 7 | 7 | 7 | this study |
| BC | MP-BA-02 | Blowhole | 50.58973 | -126.75978 | 6 | 5 | 5 | 5 | this study |
| BC | MP-BA-07 | Malcolm | 50.66542 | -126.98442 | 7 | 6 | 4 | 3 | this study |
| BC | MP-PH-01 | Storey Beach | 50.71028 | -127.41769 | 7 | 7 | 7 | 7 | this study |
| BC | MP-QS-05 | Pamphlet Cove | 50.52264 | -127.65607 | 7 | 7 | 7 | 6 | this study |
| BC | MP-QS-01 | Kains Island | 50.44589 | -128.03422 | 7 | 7 | 5 | 5 | this study |
| BC | MP-KQ-02 | SE Spring Island | 49.99919 | -127.42632 | 7 | 7 | 7 | 7 | this study |
| BC | MP-KQ-04 | Surprise Island | 50.03875 | -127.29875 | 7 | 7 | 7 | 7 | this study |
| BC | MP-KQ-06 | Rugged Point | 49.97126 | -127.25103 | 7 | 7 | 7 | 7 | this study |
| BC | MP-CS-04 | Hot Springs Cove | 49.35333 | -126.26083 | 7 | 6 | 6 | 6 | this study |
| BC | MP-CS-01 | Obstruction Island | 49.39472 | -126.08306 | 5 | 2 | 2 | 2 | this study |
| BC | MP-CS-05 | ‡ Catface | 49.24061‡ | -126.01013‡ | 7 | 6 | 5 | 4 | this study |
| BC | MP-BS-03 | Turret Dive Site | 48.89752 | -125.34417 | 7 | 7 | 5 | 5 | this study |
| BC | MP-BS-04 | Effingham Bay | 48.87885 | -125.31983 | 7 | 7 | 7 | 7 | this study |
| BC | MP-BS-06 | Mullins Island | 48.90806 | -125.29760 | 7 | 7 | 7 | 7 | this study |
| BC | MP-BS-08 | Btwn Jacques and Gibraltar | 48.91678 | -125.26225 | 7 | 7 | 6 | 6 | this study |
| BC | MP-BS-09 | Dodd Island | 48.92548 | -125.32777 | 7 | 7 | 7 | 6 | this study |
| BC | MP-BS-10 | Hand Island | 48.95360 | -125.30935 | 7 | 7 | 7 | 7 | this study |
| BC | MP-BS-21 | Tzartus Backside | 48.91873 | -125.10332 | 7 | 7 | 7 | 5 | this study |
| BC | MP-BS-27 | Nanat | 48.88055 | -125.07620 | 7 | 7 | 7 | 7 | this study |
| BC | MP-BS-26 | Danvers Danger Rocks | 48.87600 | -125.09220 | 7 | 7 | 6 | 5 | this study |
| BC | MP-BS-16 | Helby | 48.85527 | -125.16855 | 7 | 7 | 7 | 7 | this study |
| BC | MP-BS-25 | Grappler | 48.83841 | -125.13508 | 7 | 7 | 7 | 6 | this study |
| BC | MP-BS-23 | Second Beach | 48.81440 | -125.17195 | 7 | 6 | 6 | 6 | this study |
| BC | MP-BS-15 | Dodger | 48.82968 | -125.19634 | 7 | 7 | 4 | 3 | this study |
| BC | MP-BS-14 | BBNEK | 48.82727 | -125.21169 | 6 | 6 | 5 | 3 | this study |
| BC | MP-BS-13 | Skull Bay | 48.82637 | -125.22238 | 7 | 7 | 6 | 5 | this study |
| BC | MP-BS-22 | Island by Mudd Cove | 48.79347 | -125.21465 | 7 | 7 | 3 | 3 | this study |
| WA | MP-WC-01 | Tatoosh Island | 48.39361 | -124.73515 | 5 | 5 | 5 | 5 | this study |
| BC | MP-SK-01 | Ella Beach | 48.35709 | -123.74846 | 7 | 7 | 6 | 4 | this study |
| CA | MP-CAi-01 | Stillwater Cove (*integrifolia*) | 36.56506 | -121.94355 | 5 | 4 | 2 | 1 | Gonzalez et al.[19] |
| CA | MP-CAp-01 | Stillwater Cove (*pyrifera*) | 36.56506 | -121.94355 | 5 | 5 | 5 | 5 | Gonzalez et al.[19] |
| CA | MP-CA-02 | Cayucos | 35.44646 | -120.93239 | 5 | 4 | 4 | 4 | Gonzalez et al.[19] |
| CA | MP-CA-03 | Arroyo Quemado | 34.470 | -120.119 | 16 | 0 | 0 | 0 | Molano et al.[20] |
| CA | MP-CA-04 | Catalina Island | 33.462 | -118.511 | 16 | 0 | 0 | 0 | Molano et al.[20] |
| CA | MP-CA-05 | Camp Pendleton | 33.157 | -117.360 | 17 | 1 | 1 | 1 | Molano et al.[20] |
| CL | MP-CL-01 | Playa Blanca | -28.18511 | -71.16517 | 5 | 5 | 5 | 5 | Gonzalez et al.[19] |
| CL | MP-CL-02 | Punta Choros | -29.24571 | -71.46617 | 5 | 5 | 5 | 5 | Gonzalez et al.[19] |
| CL | MP-CL-03 | Totoralillo | -32.02284 | -71.50866 | 5 | 5 | 5 | 5 | Gonzalez et al.[19] |
| CL | MP-CL-04 | Las Docas | -33.13985 | -71.70822 | 5 | 5 | 3 | 3 | Gonzalez et al.[19] |
| CL | MP-CL-05 | Algorrobo | -33.36002 | -71.66873 | 5 | 5 | 5 | 5 | Gonzalez et al.[19] |
| CL | MP-CL-06 | Las Monjas | -33.49218 | -71.64249 | 5 | 4 | 4 | 3 | Gonzalez et al.[19] |
| CL | MP-CL-07 | La Boca | -33.90318 | -71.83477 | 5 | 5 | 5 | 5 | Gonzalez et al.[19] |
| CL | MP-CL-08 | Tumbes | -36.62373 | -73.09302 | 5 | 5 | 5 | 5 | Gonzalez et al.[19] |
| CL | MP-CL-09 | Chome | -36.80647 | -73.17709 | 5 | 5 | 5 | 5 | Gonzalez et al.[19] |
| CL | MP-CL-10 | Bahía Mansa | -40.58156 | -73.73820 | 5 | 5 | 5 | 5 | Gonzalez et al.[19] |
| CL | MP-CL-11 | Ancud | -41.86531 | -73.83193 | 5 | 5 | 5 | 5 | Gonzalez et al.[19] |
| CL | MP-CL-12 | Dalcahue | -42.38276 | -73.65502 | 5 | 5 | 5 | 5 | Gonzalez et al.[19] |
| AU | MP-AU-01 | Blackmans Bay | -43.01733 | 147.32939 | 1 | 1 | 1 | 1 | Iha et al.[21] |
| | | | | **Total** | **364** | **304** | **281** | **265** | |

**Table S2. Genetic differentiation and genetic distance, Related to Figure 1**
Genetic differentiation ($F_{ST}$; above diagonal) and genetic distance ($d_{XY}$, all values multiplied by $10^3$; below diagonal) between genetic clusters of (A) bull kelp and (B) giant kelp. Genetic clusters are identified according to the same colour scheme as in Figure 1, with a written description of geographic locations represented by each cluster provided for additional clarity. Cells containing $F_{ST}$ and $d_{XY}$ values are shaded as indicated in the legend for visualization purposes.

### A) Bull kelp

| **Genetic cluster** | | | | | | | |
|---|---|---|---|---|---|---|---|
| Northern British Columbia | | | 0.16 | 0.28 | 0.39 | 0.23 | 0.24 |
| West coast of Vancouver Island | | 3.34 | | 0.08 | 0.27 | 0.20 | 0.32 |
| South-central Salish Sea | | 3.26 | 2.31 | | 0.30 | 0.25 | 0.42 |
| Southern Puget Sound | | 3.29 | 2.34 | 1.85 | | 0.47 | 0.62 |
| Northern Salish Sea | | 3.12 | 2.64 | 2.28 | 2.36 | | 0.27 |
| Broughton Archipelago | | 3.08 | 2.97 | 2.74 | 2.79 | 2.15 | |

### B) Giant kelp

| **Genetic cluster** | | | | | | | | |
|---|---|---|---|---|---|---|---|---|
| North coast of British Columbia | | | 0.29 | 0.32 | 0.47 | 0.44 | 0.55 | 0.55 |
| Haida Gwaii | | 2.34 | | 0.30 | 0.45 | 0.43 | 0.55 | 0.53 |
| Central coast of British Columbia | | 2.76 | 2.81 | | 0.32 | 0.24 | 0.40 | 0.35 |
| Broughton Archipelago | | 2.75 | 2.81 | 2.60 | | 0.36 | 0.48 | 0.50 |
| Quatsino and Kyuquot Sounds | | 3.11 | 3.20 | 2.69 | 2.58 | | 0.15 | 0.20 |
| Barkley and Clayoquot Sounds | | 3.16 | 3.27 | 2.80 | 2.63 | 1.92 | | 0.26 |
| Strait of Juan de Fuca | | 3.20 | 3.33 | 2.81 | 2.70 | 2.11 | 1.88 | |

### Legend

| $F_{ST}$ | $d_{XY}$ (x $10^{-3}$) |
|---|---|
| 0.00-0.19 | 0.00-1.99 |
| 0.20-0.39 | 2.00-2.99 |
| ≥0.40 | ≥3.00 |

**Table S3. Brown algae reference genomes, Related to Figure 3**
Reference genomes of brown algae (and the outgroup *Schizocladia ischiensis*) used to assess site-specific evolutionary rates in GERP analyses. Note that genomes from Phaeoexplorer[121] (https://phaeoexplorer.sb-roscoff.fr) and PhycoCosm[122] (https://phycocosm.jgi.doe.gov) have genome versions but no accession numbers.

| Species | Order | Size (Mbp) | Source | Accession number or genome version |
|---|---|---|---|---|
| *Alaria esculenta* | Lamniariales | 487.8 | PhycoCosm | Alaria esculenta A1 |
| *Ascophyllum nodosum* | Fucales | 1,252.9 | Phaeoexplorer | Ascophyllum nodosum dioecious genome v1.0 |
| *Chordaria linearis* | Ralfsiales | 214.6 | Phaeoexplorer | genome v1.0 of Chordaria linearis ClinC8C monoicous |
| *Choristocarpus tenellus* | Discosporangiales | 164.0 | Phaeoexplorer | Choristocarpus tenellus KU2346 genome v1.0 |
| *Cladosiphon okamuranus* | Ectocarpales | 129.9 | Phaeoexplorer | Cladosiphon okamuranus Unknown S-Strain |
| *Desmarestia dudresnayi* | Desmarestiales | 441.5 | Phaeoexplorer | genome v1.0 of Desmarestia dudresnayi DdudBR16 monoicous |
| *Desmarestia herbacea* | Desmarestiales | 430.9 | Phaeoexplorer | genome v1.0 of Desmarestia herbacea DmunM male |
| *Dictyota dichotoma* | Dictyotales | 851.2 | Phaeoexplorer | genome v1.0 of Dictyota dichotoma ODC1387m male |
| *Discosporangium mesarthrocarpum* | Discosporangiales | 160.8 | Phaeoexplorer | Discosporangium mesarthrocarpum MT17_79 genome v1.0 |
| *Ectocarpus fasciculatus* | Ectocarpales | 227.6 | Phaeoexplorer | genome v1.0 of Ectocarpus fasciculatus Ec847m_Ec191_A9_m male |
| *Ectocarpus subulatus* | Ectocarpales | 242.4 | Phaeoexplorer | genome v1.0 of Ectocarpus subulatus male Bft15b |
| *Fucus distichus* | Fucales | 724.7 | Phaeoexplorer | genome v1.0 of Fucus distichus monoicous |
| *Fucus serratus* | Fucales | 1,234.4 | Phaeoexplorer | genome v1.0 of Fucus serratus male |
| *Halopteris paniculata* | Sphacelariales | 350.6 | Phaeoexplorer | genome v1.0 of Halopteris paniculata Hal_grac_a_UBK monoicous |
| *Himanthalia elongata* | Fucales | 785.5 | Phaeoexplorer | genome v1.0 of Himanthalia elongata Himel1 dioecious |
| *Laminaria digitata* | Laminariales | 447.6 | Phaeoexplorer | Laminaria digitata LdigPH10_18mv male genome v1.0 |
| *Macrocystis pyrifera* | Lamniariales | 537.5 | PhycoCosm | Macrocystis pyrifera CI_03 v1.0 |
| *Nereocystis luetkeana* | Lamniariales | 467.6 | NCBI | GCA_031213475.1 |
| *Pelvetia canaliculata* | Fucales | 570.6 | Phaeoexplorer | genome v1.0 of Pelvetia canaliculata dioecious |
| *Pleurocladia lacustris* | Ectocarpales | 220.4 | Phaeoexplorer | genome v1.0 of Pleurocladia lacustris SAG_25_93 |
| *Saccharina japonica* | Lamniariales | 548.5 | PhycoCosm | Saccharina japonica str. Ja |
| *Saccharina latissima* | Lamniariales | 531.3 | Phaeoexplorer | genome v1.0 of Saccharina latissima SLPER63f female |
| *Saccorhiza dermatodea* | Tilopteridales | 352.0 | Phaeoexplorer | genome v1.0 of Saccorhiza dermatodea SderLu1190fm monoicous |
| *Saccorhiza polyschides* | Tilopteridales | 557.5 | Phaeoexplorer | genome v1.0 of Saccorhiza polyschides SpolBR94m male |
| *Sargassum fusiforme* | Fucales | 376.2 | Phaeoexplorer | genome v1.0 of Sargassum fusiforme |
| *Schizocladia ischiensis* | Schizocladiales | 194.5 | Phaeoexplorer | genome v1.0 of Schizocladia ischiensis KU_0333 |
| *Scytosiphon promiscuus* | Ectocarpales | 193.2 | Phaeoexplorer | genome v1.0 of Scytosiphon promiscuus 110409_Ot110409_Otamoi_16_male male |
| *Sphacelaria rigidula* | Sphacelariales | 244.2 | Phaeoexplorer | Sphacelaria rigidula Sph_rig_Cal_Mo_4_1_68b_KU_MACC1312 female genome v1.0 |
| *Undaria pinnatifida* | Lamniariales | 634.5 | Phaeoexplorer | Undaria pinnatifida Kr2015 |

**Table S4. Regressions predicting genetic load in between-population crosses, Related to Figure 4**

Multiple linear regressions of variables predicting the relative difference in realized genetic load (the proportion of sites homozygous for the derived allele in deleterious allele categories) when comparing simulated between-population crosses (recipient x donor population) to within-population crosses (recipient x recipient) for (A,C) bull kelp and (B,D) giant kelp. The deleterious allele category examined was evolutionarily conserved sites in (A-B) GERP analyses and moderate- to high-impact sites in (C-D) *SnpEff* analyses. Smaller values of the response variable indicate a greater reduction in realized load in the recipient population when a cross is conducted between populations. Predictor variables include the recipient effective population size ($N_e$) (log10-scale) and the geographic distance by ocean (km) of the donor population. Adjusted two-sided $p$-values ($p_{adj}$) for each predictor variable were calculated by permuting the predictor variable 1,000 times while holding other variables constant.

**A) GERP analysis, bull kelp**

| | Estimate | Standard error | $t$ | $p_{adj}$ |
|---|---|---|---|---|
| Intercept | $-7.89 \times 10^{-2}$ | $1.73 \times 10^{-2}$ | -4.6 | |
| Recipient $\log_{10}(N_e)$ | $2.74 \times 10^{-1}$ | $5.31 \times 10^{-3}$ | 51.5 | < 0.001 |
| Donor geographic distance (km) | $-6.73 \times 10^{-4}$ | $1.14 \times 10^{-5}$ | -59.0 | < 0.001 |

**B) GERP analysis, giant kelp**

| | Estimate | Standard error | $t$ | $p_{adj}$ |
|---|---|---|---|---|
| Intercept | $5.73 \times 10^{-1}$ | $4.59 \times 10^{-2}$ | 12.5 | |
| Recipient $\log_{10}(N_e)$ | $1.23 \times 10^{-1}$ | $2.23 \times 10^{-2}$ | 5.5 | 0.188 |
| Donor geographic distance (km) | $-7.58 \times 10^{-4}$ | $1.33 \times 10^{-5}$ | -56.8 | < 0.001 |

**C) *SnpEff* analysis, bull kelp**

| | Estimate | Standard error | $t$ | $p_{adj}$ |
|---|---|---|---|---|
| Intercept | $-9.26 \times 10^{-2}$ | $1.61 \times 10^{-2}$ | -5.8 | |
| Recipient $\log_{10}(N_e)$ | $2.77 \times 10^{-1}$ | $4.93 \times 10^{-3}$ | 56.2 | < 0.001 |
| Donor geographic distance (km) | $-5.52 \times 10^{-4}$ | $1.06 \times 10^{-5}$ | -52.1 | < 0.001 |

**D) *SnpEff* analysis, giant kelp**

| | Estimate | Standard error | $t$ | $p_{adj}$ |
|---|---|---|---|---|
| Intercept | $5.64 \times 10^{-1}$ | $4.37 \times 10^{-2}$ | 12.9 | |
| Recipient $\log_{10}(N_e)$ | $1.28 \times 10^{-1}$ | $2.12 \times 10^{-2}$ | 6.1 | 0.171 |
| Donor geographic distance (km) | $-7.24 \times 10^{-4}$ | $1.27 \times 10^{-5}$ | -57.1 | < 0.001 |
